# Quorum sensing dependent repression of the *Staphylococcus aureus* transcription termination factor Rho is necessary for pathogenicity

**DOI:** 10.1101/2025.07.24.666493

**Authors:** Laura Barrientos, Marion Blum, Florence Couzon, François Vandenesch, Karen Moreau, Pascale Romby, Isabelle Caldelari

## Abstract

In *Staphylococcus aureus*, the transcription termination Rho factor has been identified as an attenuator of virulence, highlighting this universal protein as a new player in *S. aureus* pathophysiology. Moreover, Rho is required for the repression of “pervasive” transcription, which results from transcription at cryptic promoters or from transcriptional readthrough. In our study, we demonstrated that the *rho* transcript is specifically repressed by the quorum sensing-dependent RNAIII during the stationary phase of growth, allowing exotoxins to be activated at this stage. Rho-dependent RNA targets were identified using RIP-seq and differential transcriptomic analyses, revealing that several of these targets are involved in adhesion, dissemination, and the host responses. Our data provide a molecular explanation for the functions of this well-conserved termination factor, which have evolved in this human pathogen.

## INTRODUCTION

In bacteria, transcription termination is an essential step of gene expression and is also required for maintenance of genome integrity^1,2^. Transcription termination occurs through two mechanisms long considered independent from one another: intrinsic termination and factor-dependent termination. Intrinsic termination requires the formation of a terminator hairpin at the 3’ end of the nascent RNA followed by a stretch of U residues. This secondary structure alone promotes the dissociation of the transcription elongation complex. Conversely, Rho-dependent termination involves the transcription termination factor Rho, an ATP-dependent RNA-DNA helicase, that binds specific features on the nascent RNA named *rut* sites - composed of non-structured cytidine-rich regions - and releases the RNA polymerase from the DNA-RNA hybrid^3^. The Rho factor has been extensively studied in *Escherichia coli* since this protein is essential and mediates the widespread termination of about half of transcription events. In *E. coli*, the transcription elongation factor NusG has been found to stimulate Rho-dependent termination *in vitro* by directly interacting with the termination factor and accelerating RNA release at termination sites and stalled elongation complexes^4–6^. About one third of Rho-dependent terminators are NusG-dependent in *E. coli*. It is believed that the interaction of NusG with Rho induces a rapid conformational change to the active form of Rho, helping to stabilize the Rho-RNA interaction at suboptimal *rut* sites^7^. NusG is universally conserved among bacteria and its absence has global effects on gene expression and bacterial physiology, like motility in *Bacillus subtilis*^8^. In addition, NusG, together with NusA, enhance intrinsic termination at weak terminators by inducing transcriptional pausing^8,9^. More recently, it has been shown that there is an interplay between intrinsic termination and Rho to enhance termination at the 3’ ends of genes carrying inefficient intrinsic terminators in *E. coli, Mycobacterium tuberculosis*, and *B. subtilis* to prevent readthrough at these locations^10,11^.

The significance of Rho beyond simple transcription termination is also well established. This factor is important for regulation of gene expression through transcriptional polarity, riboswitch-mediated attenuation, and sRNA-mediated regulation^12,13,14^. For instance, Rho regulates the transcription of several genes, such as the *tna* operon in *E. coli* or the *mgtA* riboswitch in *Salmonella*^15^. By terminating transcription in the leader region of these operons, Rho prevents transcription of the downstream genes through a mechanism known as attenuation. Some sRNAs, such as ChiX from *Salmonella* can influence Rho-dependent termination by affecting the accessibility of the *rut* sites on target mRNAs^16^. Additionally, Rho serves as a guardian of genome integrity as it limits the expression of exogenous DNA, which could be deleterious, and recycles stalled RNA polymerase to facilitate DNA repair and other essential DNA-dependent processes for cell survival^17,18^. Most notably, Rho suppresses pervasive transcription, which frequently produces antisense (asRNAs) that can interfere with gene expression, genome stability, and replication^6^. The spurious transcription of asRNAs leads to the formation of asRNA-mRNA duplexes, which are then targeted for degradation by the double-stranded endonuclease RNase III^19,20^.

Even though Rho is well conserved among bacteria, with over 90% of genomes containing a Rho homologue, the factor is not always required for cell viability^21^. Most notably, Rho is not essential in the Gram-positive *B. subtilis* and the opportunistic human pathogen *Staphylococcus aureus,* but it accomplishes relevant biological function^21,22^. In *B. subtilis*, Rho regulates motility, sporulation, and cell fate determination^23^. Rho is also necessary for the adaption of *B. subtilis* to stationary phase, a phase characterized by physiological remodeling for bacterial adaptation to limited resources^24^. In *S. aureus*, the mechanism of action of Rho is not well studied. Rho decreases antisense transcription consistent with its conserved role in suppressing pervasive transcription^25^. Furthermore, the *rho* mutant appears more virulent than its cognate WT strain in a murine model of bacteremia showing that Rho acts as a suppressor of virulence^26^. This effect might be due to the deregulation of the SaeRS regulon, which is induced in an *S. aureus rho* deletion mutant^26^. The Sae two-component system (TCS) responds to several stimuli, especially induced by the host defenses, and controls the expression of multiple virulence genes encoding toxins, adhesins, and immune evasion proteins^27^. This system is composed of four genes: the sensor histidine kinase SaeS, the response regulator SaeR and two auxiliary proteins, SaeP and SaeQ, necessary for the activation of the phosphatase activity of SaeS^27^. Interestingly, the activation of the Sae regulon was shown to be directly dependent on the transcription termination activity of Rho, since its inhibition by bicyclomycin, an antibiotic that specifically blocks ATPase activity of Rho, has the same virulence-inducing effect^26^.

The physiological significance of Rho in different bacterial species supposes the existence of tight regulatory mechanisms to ensure the correct expression of this protein. For instance, *E. coli rho* is autogenously regulated by transcriptional attenuation of the *rho* mRNA, leading to premature termination^28,29,30^. Conversely, the sRNA SraL binds to the 5’UTR of the *rho* mRNA and counteracts premature transcription termination by competing with Rho for the *rut* sites^31^. Another layer of regulation has been recently described, where Rho can be transiently inactivated upon stress by undergoing oligomerization^32^. This sequestration mechanism is reversible and allows modulation of Rho activity during stress conditions, where the latter could be detrimental^32^. Similarly, in the Gram-negative *Acinetobacter baumanii*, where *rho* is essential, Rho levels are maintained constant but its activity is modulated by external factors to decrease Rho-dependent termination, promoting a switch between virulent and avirulent states^33^. Similarly, in *B. subtilis*, Rho amounts remain low^34^, and both *rho* mRNA and protein levels decrease upon entry intro stationary phase, suggesting the existence of regulatory pathways controlling *rho* expression^24^. In *S. aureus*, the level of *rho* mRNA is at its lowest during stationary phase, indicating that the *rho* gene regulation is crucial once the bacterial population grows, when significant physiological remodeling takes place and exotoxin activation occurs^25^.

Regulation of virulence factors in *S. aureus* involves complex regulatory networks including TCS, transcription factors, and regulatory RNAs (sRNAs). The accessory gene regulator (Agr) quorum-sensing system responds to cellular density and is a global regulator of virulence^35^. The system senses an auto-inducing peptide (AIP) and only when its extracellular concentration reaches a certain threshold, the system is activated. The *agr* locus is composed of two divergent operons initiating the transcription of divergent transcripts named RNAII and RNAIII. The RNAII mRNA encompasses the four *agrBDCA* genes that encode the TCS machinery, AgrC being the sensor and AgrA, the response regulator^35^. One of the main effector of the system is RNAIII, which is a 514 nt-long bifunctional sRNA that encodes ∂-hemolysin (*hld*)^36^. RNAIII post-transcriptionally regulates many mRNA targets by direct RNA-RNA interactions, affecting stability and/or translation of the target mRNAs^37,38,39^. For instance, it activates α-hemolysin (Hla) synthesis by favoring its mRNA translation and represses the translation of other mRNAs such as *rot*, encoding the transcriptional repressor of exotoxins, or *spa*, the mRNA encoding the immunoglobin binding protein A^40,41^. All-in-one, RNAIII coordinates the switch from the adhesion mode of early infection to a disseminative state necessary during later stage of infection.

As mentioned above, in *S. aureus, rho* mRNA levels decrease during stationary phase when the expression of exotoxins is activated. Rho has been described as a suppressor of this phenomenon by inhibiting the expression of the Sae regulon during bacterial proliferation. These observations suggest the existence of fine-tuning mechanisms for the levels of Rho to ensure proper virulence gene expression. In this work, we show that the *rho* gene undergoes post-transcriptional regulation during stationary phase, mainly by RNAIII to allow its repression. We also identified RNA targets of Rho, highlighting its direct involvement in repressing the *saePQRS* locus and virulence genes. Our findings place Rho within the main regulatory network of virulence in *S. aureus* and add transcription termination to the vast repertoire of sophisticated mechanisms this human pathogen has developed to adapt and colonize its host.

## RESULTS

### Expression of *rho* mRNA is quorum-sensing dependent

In *S. aureus,* Rho is most likely involved in the regulation of virulence pathways under the control of the quorum sensing system ^25^. Therefore, we first analysed the steady-state levels of *rho* mRNA during cell growth. The expression of the *rho* mRNA was monitored at different time points during growth of HG001 WT strain (Fig. 1A). Culture samples were taken at early exponential (2 h of growth), mid exponential (4 h) and early stationary phases (6 h) for RNA extraction and visualization on Northern blot. We observed an accumulation of the *rho* mRNA during early exponential phase, followed by a strong decline during stationary phase (Fig. 1A).

**Figure 1.**
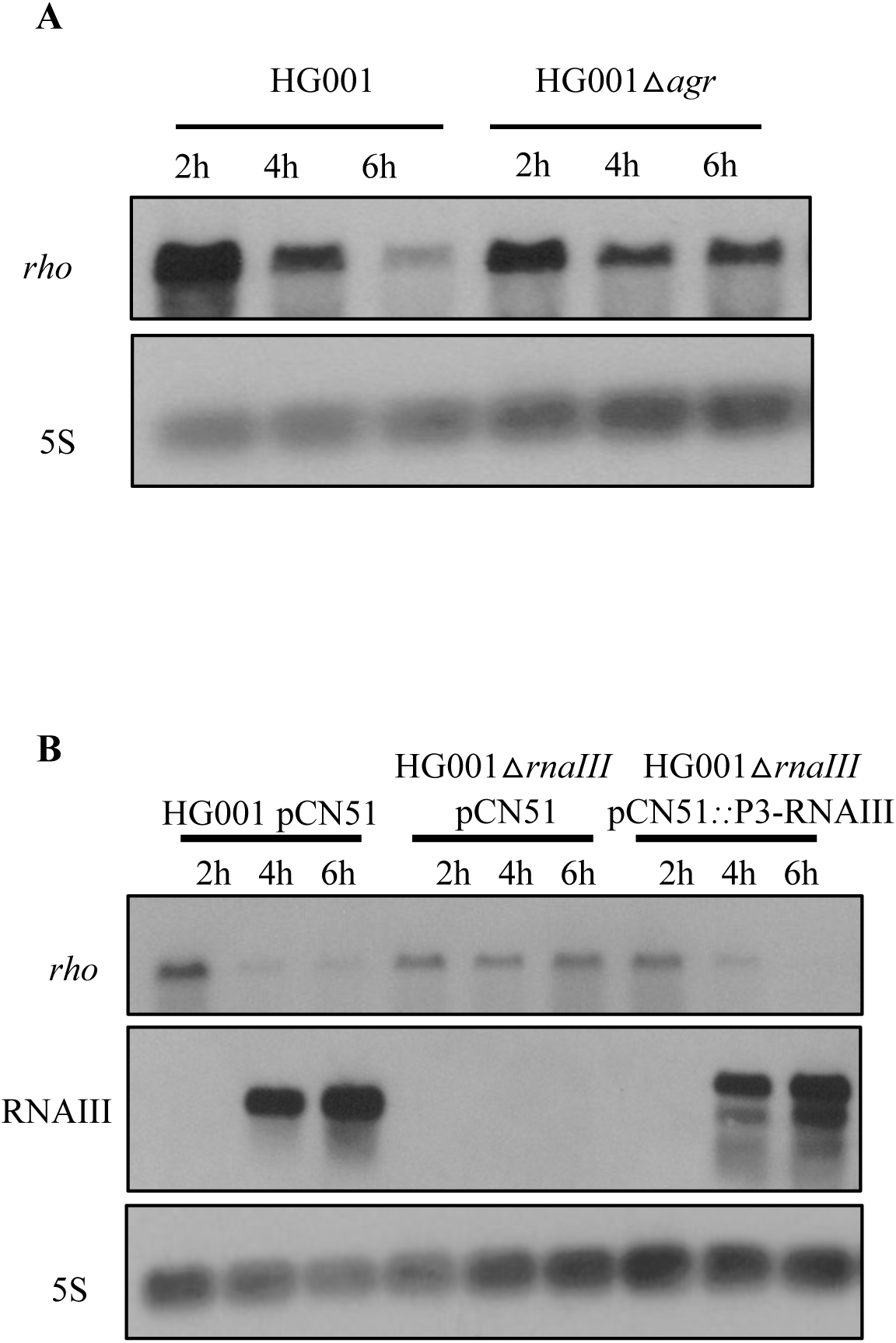
*rho* mRNA is regulated by the quorum sensing *agr* system. **A.** Northern blot analysis of *rho* mRNA in HG001 WT and HG001 △*agr* strains. Total RNA was extracted at 2, 4, and 6 h of growth in BHI medium at 37°C and loaded on a 1% agarose gel. DIG-labeled RNA probes against *rho* and the loading control 5S rRNA (5S) were used. The 5S rRNA was probed on the same membrane. **B**. Northern blot analysis of *rho* mRNA and RNAIII in HG001 pCN51, △*rnaIII* pCN51 and △*rnaIII* pCN51::RNAIII strains. Total RNA was extracted at 2, 4 and 6 h of growth in BHI medium supplemented with 10 µg/mL erythromycin and loaded on a 1% agarose gel. DIG-labeled RNA probes against *rho,* RNAIII, and the loading control 5S rRNA (5S) were used. RNAIII and 5S rRNA were probed from the same gel as *rho* mRNA.

Given that *rho* expression of is growth-phase dependent, we examined the levels of *rho* mRNA in various HG001 mutant strains, each lacking one of the two-component systems (TCS) that detect and connect environmental changes to physiological responses (Fig. S1)^42^. Among the tested TCS, we found that the *agr* mutant strain alleviates the repression of *rho* at the late stationary phase of growth (Fig. S1, Fig. 1A). Since the regulatory bi-functional RNAIII is one of the main intracellular effectors of the *agr* system, we analysed the yields of *rho* mRNA in the deletion mutant Δ*rnaIII* HG001 strain, as well as in the same mutant strain complemented with a plasmid expressing RNAIII. We found that in the Δ*rnaIII* mutant strain, expression of *rho* becomes constitutive while the phase-dependent repression of *rho* was restored in the complemented strain expressing RNAIII similar to what is observed in the WT HG001 strain (Fig. 1B). Note that RNAIII transcribed from the plasmid is slightly longer than the chromosomal copy, most probably due to the weakness of its terminator, leading to the usage of the terminator sequence encoded on the plasmid. However, this longer RNAIII is still able to regulate key virulence factors, as the activation of *hla* was restored in the complemented mutant strain (Fig. S2). The RNAIII-dependent repression of *rho* was further confirmed in *S. aureus* RN6390 WT strain, in which RNAIII expression is stronger than in HG001 (Fig. S3).

All in all, our results show that the quorum sensing *agr* system, through its main effector RNAIII, represses *rho* during stationary phase.

### RNAIII binds to *rho* mRNA at multiple sites

RNAIII is a long, highly structured sRNA with 14 hairpins and contains the coding region for the ∂-hemolysin gene (*hld*)^43^. To assess the existence of a *rho*-RNAIII interaction *in vivo*, we first applied the MAPS technique (MS2 affinity purification coupled with RNA sequencing) which was previously used to determine the RNA targetome of several staphylococcal sRNAs^44,45,46^. Briefly, RNAIII was tagged with an MS2 tag at its 5’ end and expressed in the Δ*rnaIII* mutant strain from a plasmid under the control of its own P3 promoter. This tagged version of RNAIII exhibited very similar steady-state levels as compared to the WT RNAIII (Fig. S4). After purification by affinity chromatography, RNAs from each fraction were extracted and analyzed by Northern blot using specific probes against RNAIII and *rho* mRNA. The MS2-RNAIII was specifically retained by the MS2-MBP fusion protein and the *rho* mRNA was significantly enriched in the elution fractions of MS2-RNAIII (Fig. 2).

**Figure 2.**
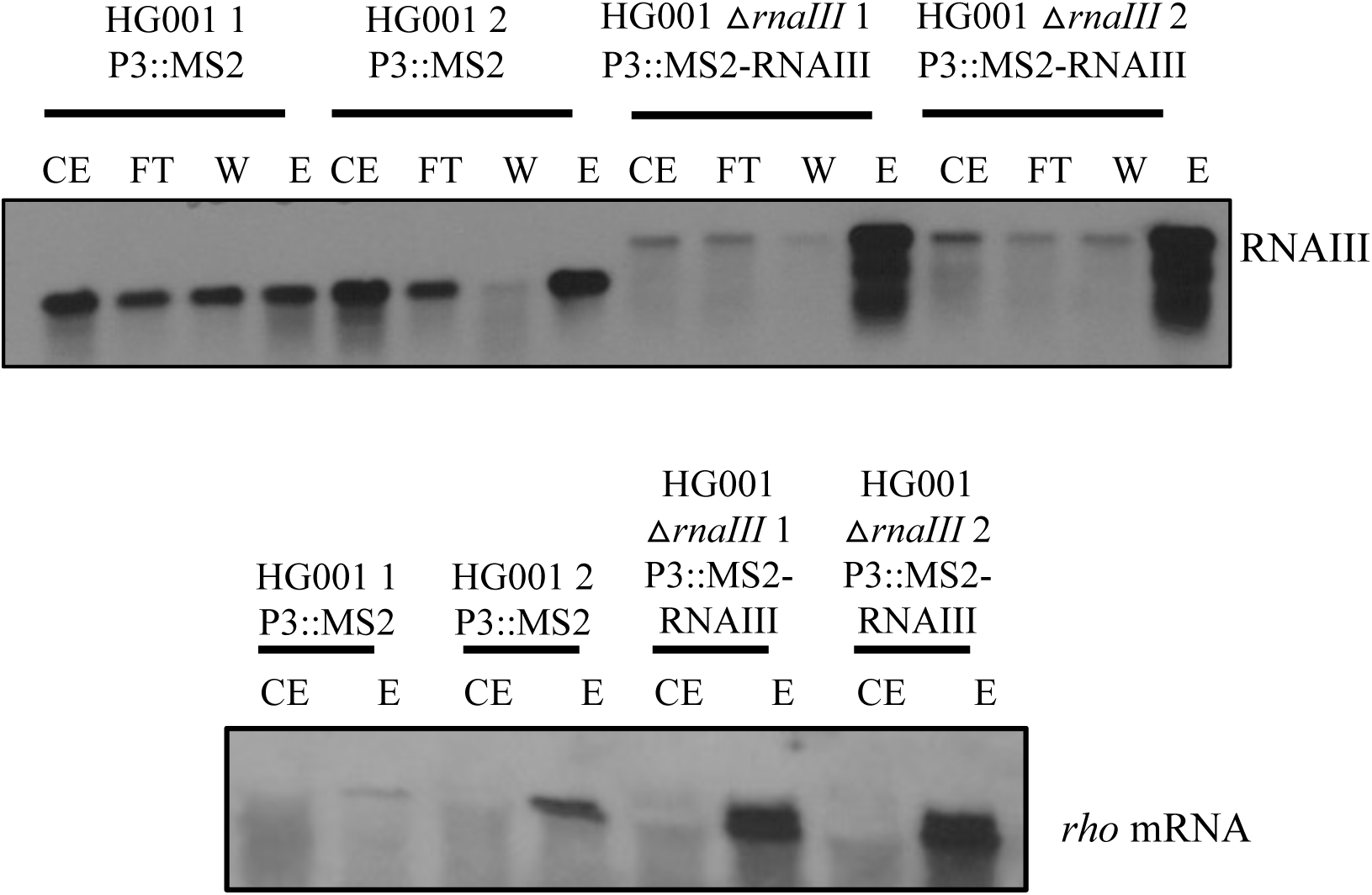
*rho* mRNA was pooled down with RNAIII *in vivo*. MS2-affinity purification of MS2-RNAIII was performed in a △*rnaIII* background using HG001 strain expressing the MS2-tag alone as a control. HG001 pCN51::P3 MS2 and △*rnaIII* pCN51::P3 MS2-RNAIII strains were grown in BHI medium supplemented with 10 µg/µL erythromycin for 6 h. Cells were harvested, lysed and the supernatant was subjected to affinity purification. RNAs from each fraction (CE: crude extract, FT: flowthrough, W: wash, E: elution) were precipitated and loaded on two separate 1% agaroses gel to perform Northern blot. Specific DIG-labeled RNA probes against RNAIII and *rho* mRNA were used. The data of two different replicates are shown (1, 2).

*In silico* predictions of base-pairing interactions between RNAIII and *rho* mRNA were then performed using IntaRNA^47–50^. The CU-rich unpaired region of hairpin 9 of RNAIII was predicted to interact with two distant regions of the coding sequence of the *rho* mRNA. These two regions included either nucleotides +46 to +82 (Fig. 3A) or nucleotides +1167 to +1224 near the 3’-end of *rho* mRNA (Fig. S5A). To determine whether RNAIII and *rho* mRNA can form a stable complex, we conducted gel retardation assays using *in vitro* transcribed *rho* mRNA and the full length RNAIII labeled at its 5’ end. Two different fragments of *rho* mRNA were used respectively, the 5’-end containing nucleotides 1 to 468 and the 3’ end of the mRNA encompassing nucleotides 965 to 1417. A stable interaction with the 5’ part of *rho* could be visualized *in vitro* (Fig. 3B) and quantification indicated that the dissociation constant (Kd) is around 150 nM. Unexpectedly, the deletion of hairpin 9 of RNAIII (RNAIII △H9) had only a minor effect on the binding to *rho* mRNA in gel retardation assay (Fig. 4A) suggesting that there is an alternative binding site on RNAIII (Fig. 3A).

**Figure 3.**
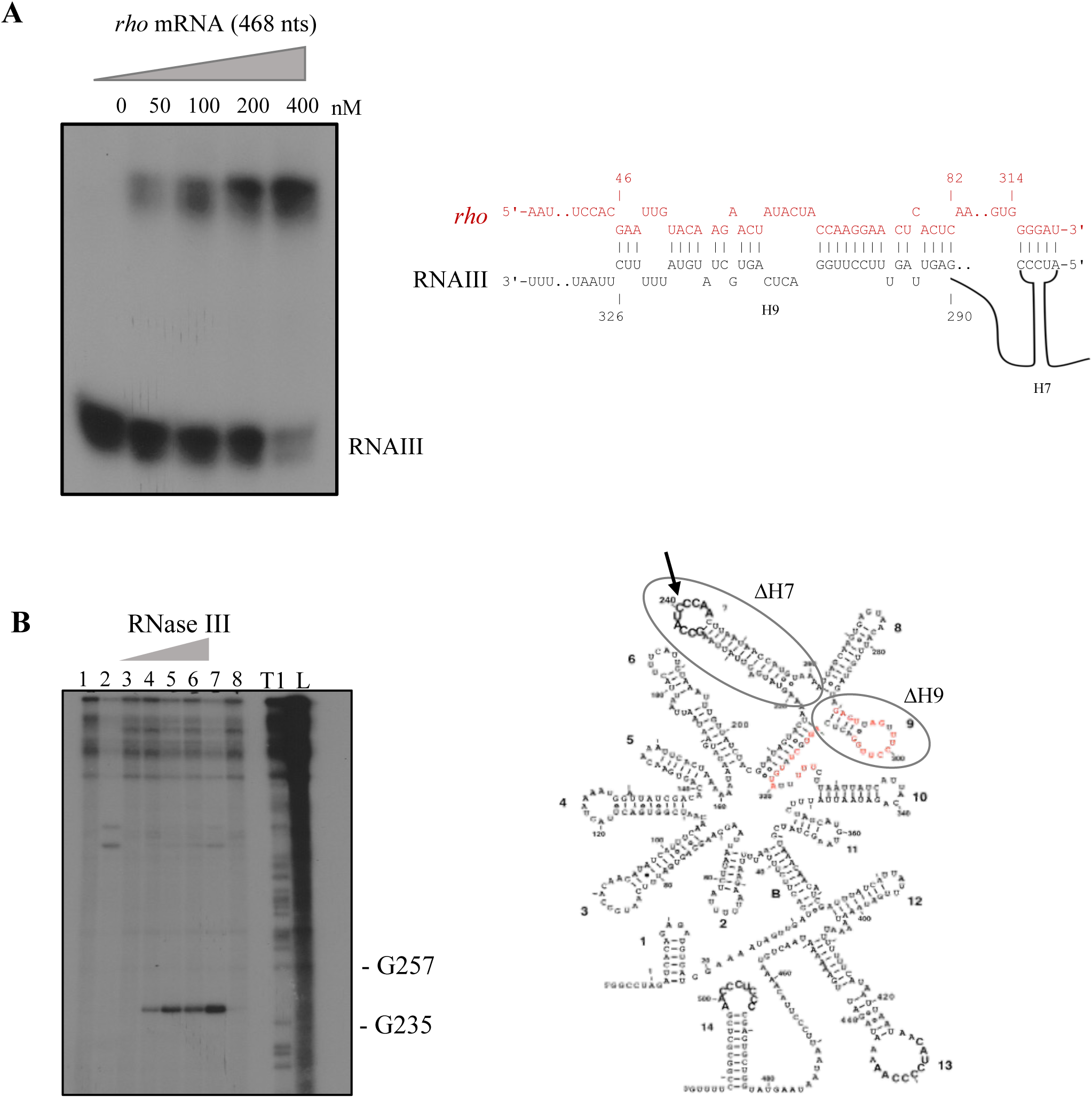
RNAIII binds to *rho* mRNA *in vitro*. **A**. Electrophoretic mobility shift assay shows the formation of the complex between 5’-end radiolabeled RNAIII and a 468 nt-long fragment of the *rho* mRNA from the 5’ end comprising the region predicted to interact with RNAIII. The 5’-end radiolabeled RNAIII was incubated with increasing concentrations of cold rho mRNA (50 to 400 nM) (left side). On the right side is the predicted interactions in hairpins 7 and 9. **B**. The RNAIII-*rho* duplex is cleaved by RNase III *in vitro.* (Left) 5’-end radiolabeled RNAIII is incubated with the *rho* mRNA fragment (468 nts from the 5’ end) without or with increasing concentrations of purified *S. aureus* RNase III. Lane 1: incubation control of RNAIII in the absence of RNase III; Lane 2: RNase III (200 nM) cleavage assay of RNAIII; Lane 3: incubation control of RNAIII with 400 nM *rho* in the absence of RNase III; Lanes 4-7: hydrolysis of the RNAIII-*rho* complex in the presence of increasing concentrations of RNase III (25 nM, 50 nM, 100 nM and 200 nM). Reactions were performed in the presence of Mg^2+^ required for RNase III activity. Lane 8: reaction control using Ca^2+^ instead of Mg^2+^ to inhibit RNase III activity. Lanes T1, L: RNase T1 and alkaline ladders, respectively, under denaturing conditions (left side). The secondary structure model of RNAIII is from Benito, *et al*^43^ showing deletions of hairpin 7 (ΔH7) and hairpin 9 (ΔH9) and the identified strong RNase III cleavage in hairpin loop H7 (arrow) (right side). The RNAIII regions predicted by IntaRNA complementarity to *rho* are colored in red.

**Figure 4:**
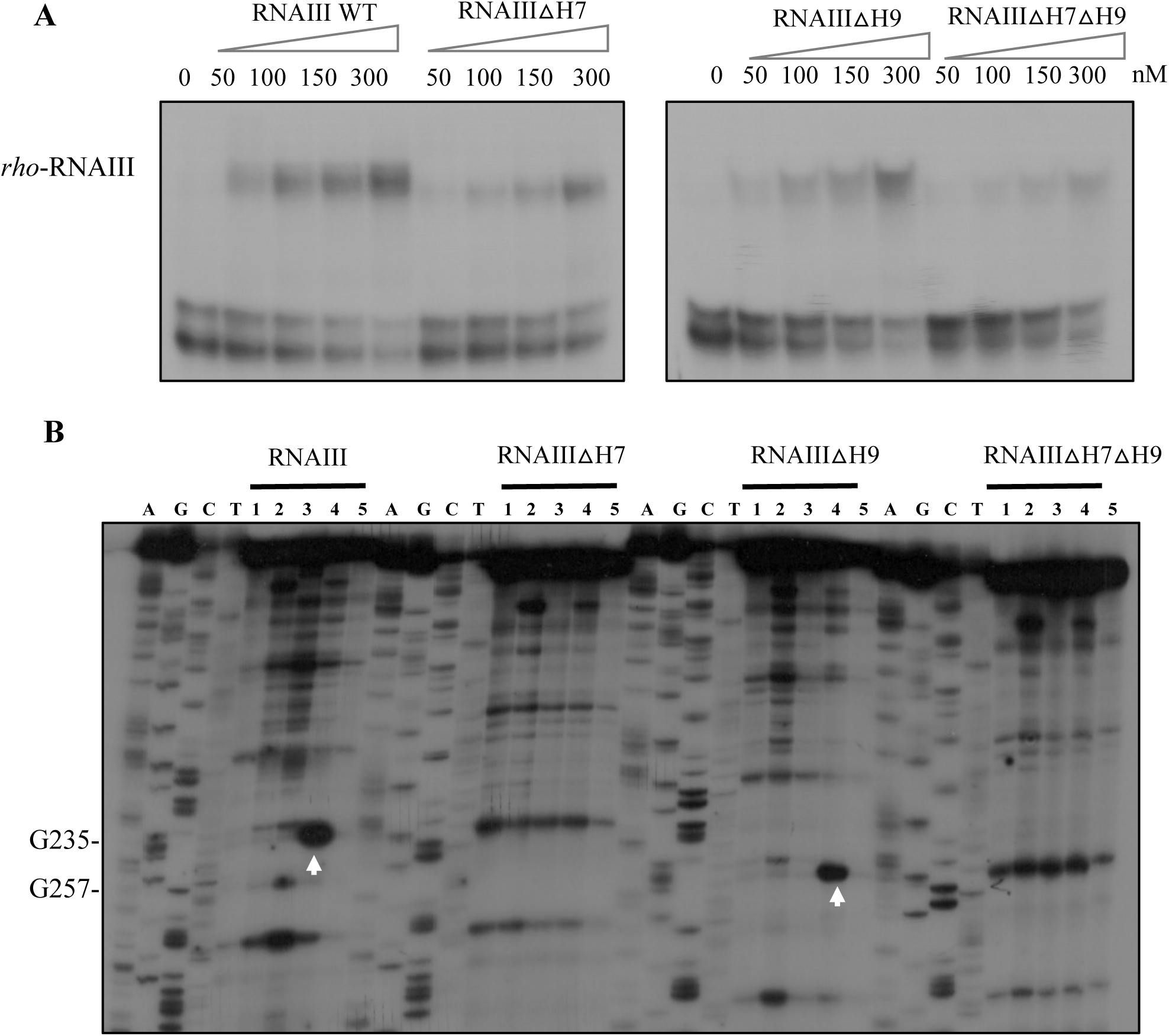
RNAIII binds to rho mRNA at multiple sites. **A**. Electrophoretic mobility shift assays show the formation of complexes between RNAIII and the 5’-end radiolabeled 468 nt-long fragment of the *rho* mRNA. Several truncated versions of RNAIII were deleted of either one hairpin H7 (ΔH7) or H9 (ΔH9), or the two hairpins H7 and H9 (ΔH7ΔH9). The 5’-end radiolabeled *rho* was incubated with increasing concentrations of cold RNAIII (50 to 300 nM). **B**. Effect of deletion of H7 and H9 of RNAIII on RNaseIII hydrolysis in the presence of *rho* mRNA. *In vitro* transcribed RNAIII, RNAIII△H7, RNAIII△H9 and RNAIII△H7△H9 (200 nM) were incubated with *rho* mRNA with or without 200 nM of RNase III in the presence of Mg^2+^ or Ca^2+^. Lane 1: incubation control of RNAIII in the absence of RNase III; Lane 2: Hydrolysis of RNAIII in the presence of 200 nM RNase III; Lane 3: incubation control of RNAIII with 400 nM *rho* in the absence of RNase III; 4: Hydrolysis of RNAIII bound to *rho* mRNA (400 nM) in the presence of RNase III (200 nM); Lane 5: Reaction control using Ca^2+^ instead of Mg^2+^ to inhibit RNase III activity; Lanes A, G, C, TA are DNA sequencing ladders. White arrows denote the major RNase III cleavage in *rho* mRNA bound to RNAIII.

RNase III has previously been shown to specifically cleave various RNAIII-mRNA complexes, making it a useful tool for identifying the targeted C-rich motifs^51^. We thus tested its ability to cleave the RNAIII-*rho* mRNA complex *in vitro* (Fig. 3B). The hybrid formed *in vitro* was incubated with increasing concentrations of purified *S. aureus* RNase III. Appropriate incubation controls were done and the cleavage positions were assigned using 5’ end-labeled RNAIII subjected to RNase T1 cleavage and alkaline hydrolysis. A strong RNase III cleavage was detected in the hairpin loop 7 of RNAIII at the C residues while no cleavage was observed in hairpin 9 (Fig. 3B). This cleavage is highly specific, as substituting Mg^2+^ with Ca^2+^ ions—an action that inhibits enzyme activity—abolishes the cleavage signal (lane 8 in Fig. 3B). This strongly indicates that hairpin 7 is also part of the *rho* binding site, which contains several G-rich motifs complementary to the C-rich sequence of hairpin loop 7 of RNAIII (Fig. 3B).

To evaluate the contribution of hairpins 7 and 9 in the formation of a stable complex with *rho* mRNA, we have synthesized RNAIII mutants lacking either hairpin 9 (RNAIII △H9), hairpin 7 (RNAIII △H7) or both hairpins (RNAIII △H7△H9). We then performed gel retardation assays using the 5’-end labeled *rho* mRNA incubated with increasing concentrations of each RNAIII variants. Deleting one hairpin had only a minor effect on *rho* binding compared to the WT RNAIII (Fig. 4A). However, deletion of both hairpins significantly reduced the interaction with *rho* mRNA, although some weak binding was still detectable. We then assessed how these mutations in RNAIII affected the RNase III cleavage patterns. Each RNAIII mutant were incubated with or without *rho* mRNA, and the formed complexes were subjected to RNase III cleavage. To detect the cuts, reverse transcription was performed using a 5’ end radiolabeled primer complementary to RNAIII (Fig. 4B). Deletion of hairpin 9 did not change the cleavage observed in hairpin 7 of RNAIII bound to *rho* (Fig. 4B), suggesting that hairpins 7 and 9 have independent binding sites on *rho*.

All these data point to a direct interaction of RNAIII with the *rho* mRNA and that RNAIII interacts at several sites of *rho* mRNA.

### Binding of RNAIII to *rho* mRNA inhibits its translation

To determine whether RNAIII affects translation of *rho* mRNA, we first examined if RNAIII fixation impacts the formation of a simplified initiation ribosomal complex on *rho* mRNA using toe-printing assays (Fig. 5A). Binding of the initiator tRNA^fMet^ and of the *S. aureus* 30S subunits to *rho* mRNA causes a typical pause of the reverse transcriptase at position +16, the so-called toe-print signal, which strongly decreased even at the lowest concentration of RNAIII. This result indicates that RNAIII hinders the ribosome binding site (Fig. 5A). In contrast, the addition of RNAIII Δ7Δ9 only reduced the toe-print signal at its highest concentration (150 nM). Interestingly, we observed two strong reverse transcriptase stops at G+73 and G+72 of *rho* mRNA caused by RNAIII binding in the absence or presence of the 30S subunit, which were not detected when RNAIII Δ7Δ9 was added in the assays. This observation correlates well with the predicted base-pairing interactions between the coding region of *rho* mRNA (region +46 to +82 from the AUG codon) and hairpin 9 of RNAIII.

**Figure 5.**
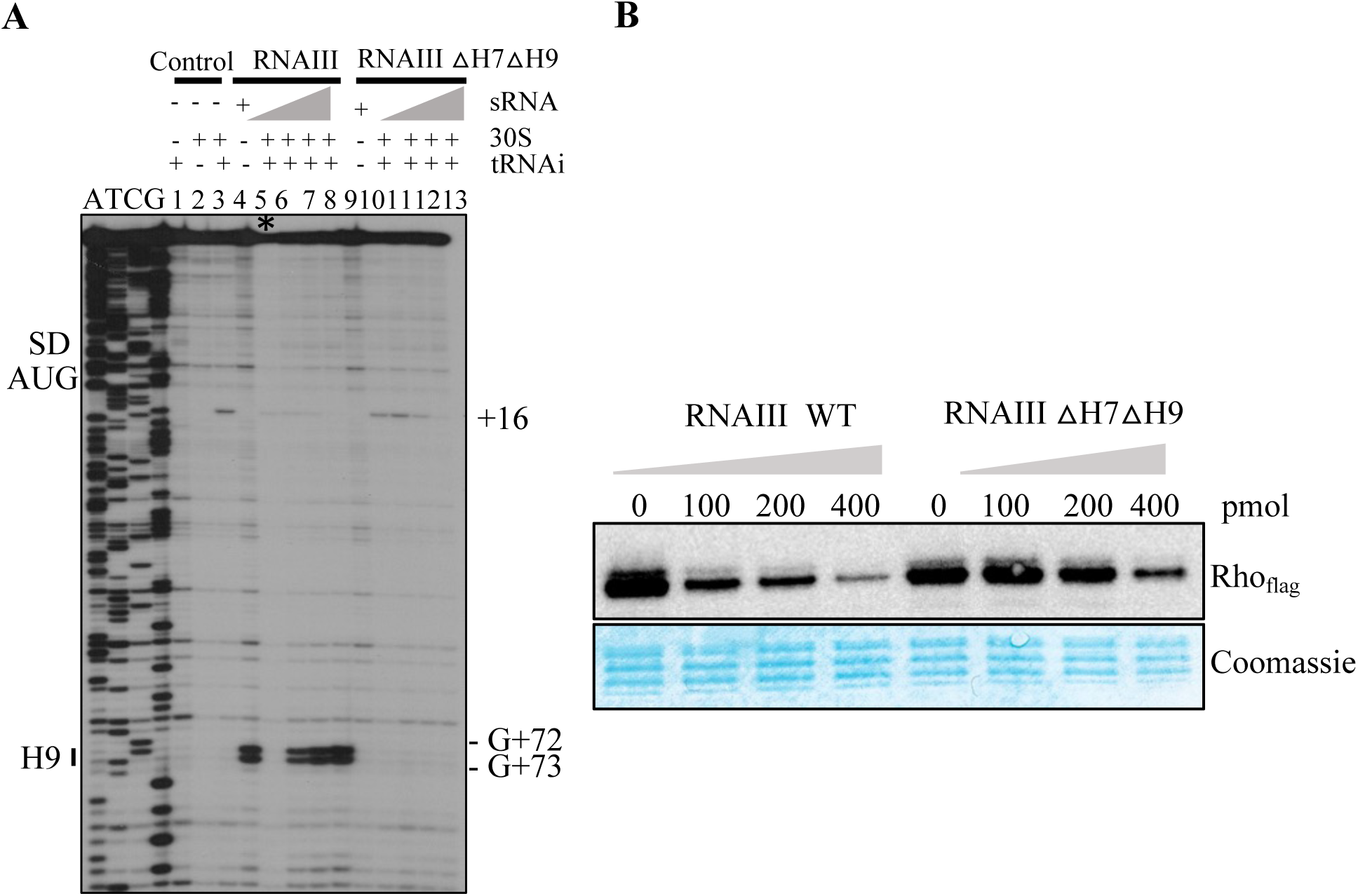
RNAIII affects translation of the *rho* mRNA *in vitro*. **A.** Toe-print assays showing the binding of *S. aureus* ribosomes on *rho* mRNA in the absence or presence of increasing concentrations of RNAIII WT or RNAIII△H7△H9. Lanes 1 & 2, incubation controls of mRNA alone with or without 30S subunit respectively; lanes 3 & 5, toe-printing assays on *rho* mRNA in the absence of RNAIII; lane 4, incubation control of *rho* mRNA with RNAIII WT (150 nM); lanes 6-8, toe-printing in the presence of increasing concentrations of RNAIII (40, 80, 150 nM); lane 9, incubation control of *rho* mRNA with RNAIII△H7△H9 (150 nM); lane 10, toe-printing assay on *rho* mRNA in the absence of RNAIIIΔH7ΔH9; lanes 10-13, toe-printing assays performed with increasing concentrations of RNAIII△H7△H9 (40, 80, 150 nM). Lanes A, T, C, G : DNA sequencing ladders. The Shine and Dalgarno (SD) sequence, the start site of translation (ATG), the toe-printing signal (+16), and the reverse transcription stops caused by RNAIII binding (H9) are indicated. (*) Note that less amount of radioactivity material was present in lane 5. **B.** *In vitro* translation assays of *rho-flag* mRNA in the absence or presence of increasing amounts of RNAIII WT or RNAIII△H7△H9 in the presence of *S. aureus* 70S ribosomes. Reactions were loaded on a 12% polyacrylamide-SDS gel. For the Western blot, membranes were incubated with anti-FLAG mouse antibodies followed by the incubation with a goat anti-mouse peroxidase (HRP) antibody. Coomassie staining of the membrane was used as loading control.

We then performed *in vitro* translation assays using *S. aureus* ribosomes and the first 468 nucleotides of *rho* mRNA with a Flag-tag sequence at the 3’ end. Western analysis with anti-FLAG mouse antibodies revealed the presence of the Rho protein (Fig. 5B). The addition of increasing concentrations of RNAIII significantly decreased Rho protein levels, while the addition of the mutant RNAIII Δ7Δ9 only slightly inhibited *rho* translation at the highest concentration (Fig. 5B).

These data strongly suggested that binding of RNAIII to *rho* mRNA represses its translation *in vitro*.

### Identification of Rho-dependent pathways determined by RNAseq

Previous tiling-array studies have shown that *rho* deletion lead to an overall increase levels of antisense RNAs and transcripts belonging to the SaeRS regulon^26^. To get a genome-wide analysis of Rho functions, we constructed a deletion mutant of *rho* in HG001 by homologous recombination. We then performed RNAseq on total RNA extracts prepared from cultures of HG001 WT and the isogenic HG001*△rho* mutant grown in BHI medium at 37°C until early exponential phase, when *rho* mRNA is mostly expressed. The experiments were done in duplicates with high reproducibility. We have considered a gene to be regulated by Rho when a two-fold expression change was observed between both strains with adjusted p-values (padj) below a 0.05 threshold (Table S1). Most differentially expressed genes in the *△rho* mutant are depicted in the volcano plot (Fig. 6A) allowing the identification of different gene categories under the regulation of Rho. The data revealed significant changes in expression consisting primarily in the upregulation of virulence factors, antisense transcripts, and prophage genes in the Δ*rho* mutant strain (Table S1A). Hence, the deletion of *rho* causes pervasive and antisense transcripts while differential regulation was observed as the result of readthrough at putative Rho-dependent sites sometimes associated with Rho-independent terminators.

**Figure 6.**
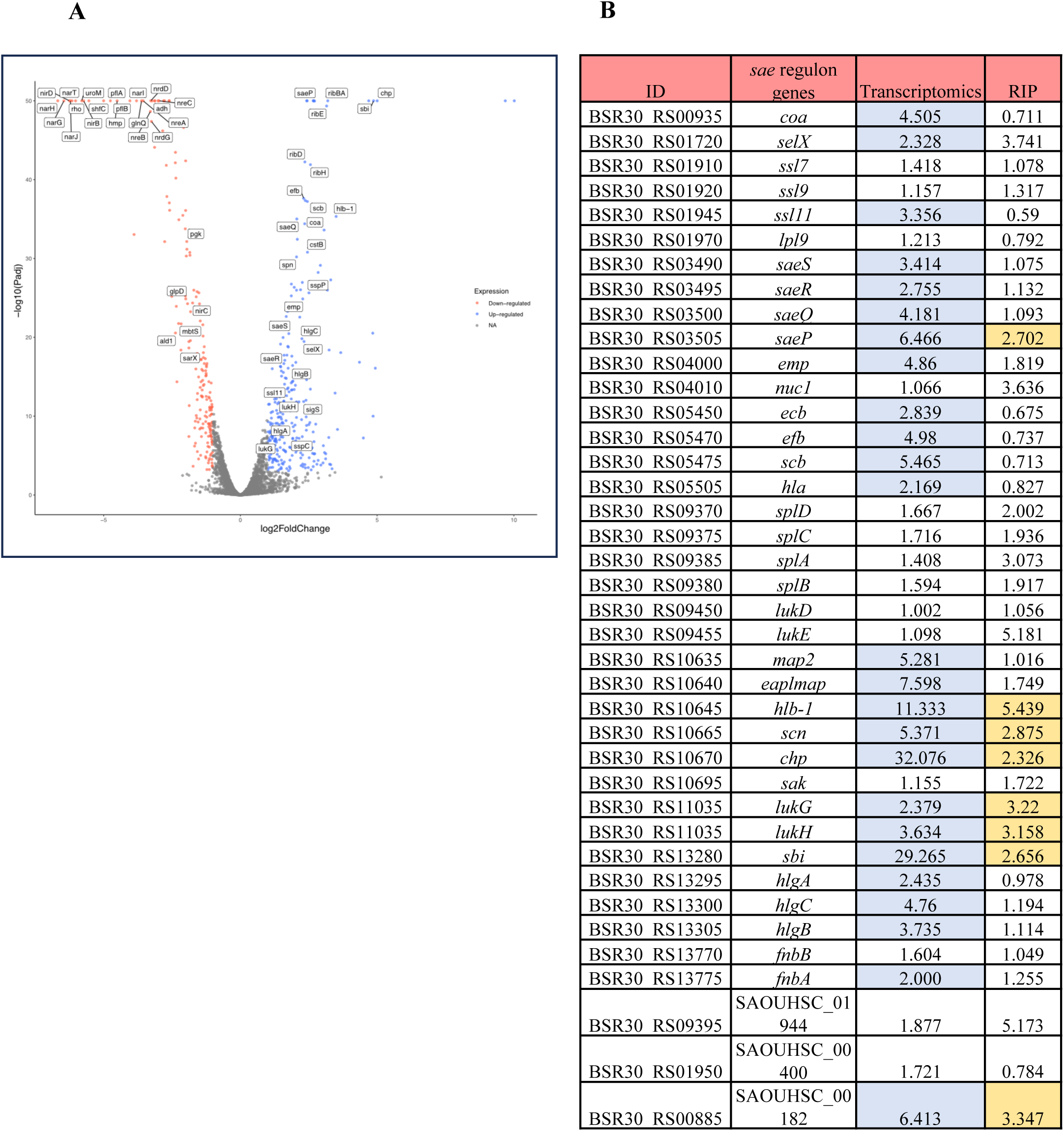
Rho regulates the expression of numerous virulence genes. **A.** Volcano plot of differentially expressed genes between HG001 WT and HG001*△rho* strains. RNAseq analysis was performed on purified RNAs from duplicates of each strain grown in BHI medium at 37°C until exponential phase. Each point represents one gene. Depicted in blue are strongly upregulated genes and in red downregulated genes in the *△rho* mutant strain. **B.** Comparison of the differential transcriptomics and RIP-Seq data for the genes belonging to the SaeRS regulon. For the RIPseq, the endogenously 3xFLAG-tagged Rho protein was immunoprecipitated from cultures grown in BHI medium at 37°C until exponential phase, and co-immunoprecipitated RNAs were isolated and sequenced. In blue are genes presenting FC>2 with padj<0.05 in the RNAseq data and in yellow the genes with FC>2 with padj<0.05 in the RIPseq data.

Several genes were downregulated upon deletion of *rho* and concomitantly increased levels of antisense RNAs were observed (Table S1B). This is particularly the cases of genes associated to prophages and all the genes involved in nitrate respiration (*narG*, *narH*, *narI* and *narJ*), transport (*narT*), and reduction (*nirB*, *nirD*), and the *nreABC* operon encoding TCS involved in the regulation of nitrate/nitrite reduction (Fig. S6A). Another strong enhanced RNA antisense was observed in the Δ*rho* mutant strain with a concomitant decreased of the yield of the corresponding operon *mbtS*, BSR_RS14940, and BSR_RS14945 (Fig. S6B). Interestingly, MbtS is a membrane bound transcriptional factor regulating the cysteine protease staphopain A and serine proteases SplA-F, all involved in host defenses^52^. In agreement with this observation, the mRNAs encoding the staphopain A (SspP) and its inhibitor SspC were upregulated (Table S1A). Other genes involved in anaerobic metabolism such as *hmp* encoding the flavohemoprotein, *ldh1* or *ldhD1* encoding lactate dehydrogenases, *adh* encoding alcohol dehydrogenase, *pflA* and *pflB* encoding respectively the pyruvate formate lyase and formate acetyl transferase enzymes, and *srrAB* encoding the TCS SrrAB (staphylococcal respiratory response), which promotes resistance to nitrosative stress and hypoxia^53^, are repressed in the Δ*rho* mutant without the obvious presence of antisense RNAs (Table S1B). Hence, downregulation of *mbtS* and *srrAB* mRNAs in the Δ*rho* mutant strain is expected to induce indirect regulatory effects.

In the Δ*rho* strain, many upregulated genes encode virulence factors. Notably, a large proportion of these genes are part of the *sae* regulon, including the *saeP* and *saeQRS* transcripts. Specifically, 25 out of 39 SaeRS-dependent genes are significantly upregulated in the Δ*rho* mutant, including major exotoxins such as HlgA, HlgB, HlgC, Hla, and LukGH (Fig. 6B). Within the four genes in the *saePQRS* operon, *saeP* is the most upregulated (Fig. 6B, S7). Among the most upregulated genes from the *sae* regulon, *chp* encodes the chemotaxis inhibitory protein^54^, *sbi* a major immunoglobulin-binding protein^55^, and *hlb-1* the non-pore forming hemolysin beta^56^ (Fig. S8, S9). For *hlb-1* and *sbi*, the regulation is most probably occurring at the transcriptional level. For the *chp* gene, a readthrough generates a long transcript antisense to the downstream genes encoding a putative phage amidase (BSR30_RS10675) and the staphylokinase SAK (BSR30_RS10680) in the Δ*rho* strain (Fig. S9). Surprisingly, this antisense RNA has no significant effect on the level of the complementary mRNA (Fig. S9). Noteworthy, a Rho-independent terminator structures are predicted at the 3’-end of the *chp* gene and at the same position on the opposite strand at the 3’-end of BSR30_RS10675, creating “mirror” terminator hairpins on both strands. Similarly, in the *Δrho* mutant, a readthrough transcript was also produced from the BSR30_RS10675 gene on the opposite strand and terminated at the Rho-independent terminator of the *scn* gene (Fig. S9). This large transcript generated an antisense RNA to the *chp* gene comprising the sRNA SprD, a translational repressor of *sbi* mRNA (Fig. S9)^57,58^.

The riboflavin synthesis operon comprising the genes *ribD*, *ribE ribAB* and *ribH* under the control of the transcriptional FMN riboswitch was upregulated in the Δ*rho* mutant (Fig. S10, Table S1A). Higher transcription levels of Teg74 and the downstream *rib* genes might result from transcriptional readthrough at Teg74 when Rho is absent (Fig. S10).

Overall, differential RNA-seq provided a global view of the impact of Δ*rho* deletion, revealing several Rho-dependent pathways linked to metabolism and virulence.

### The RNA interactome of Rho studied by RIP-Seq

In addition to the RNA-seq approach, we unveiled the RNA interactome of Rho by performing RIP-seq (RNA immunoprecipitation followed by RNA-seq). In this experiment, Rho was co-immunoprecipitated along with its RNA partners, which were then identified by deep sequencing. To perform RIP-seq, we constructed a strain expressing a chromosomally encoded tagged version of the Rho protein using homologous recombination. A triple FLAG-tag was added to the C-terminal of the protein. The steady-state levels of the modified *rho* mRNA were like those of the WT *rho* mRNA (Fig. S11A) and the Rho-3xFLAG protein was well expressed (Fig. S11B). The Rho-3xFLAG was immunoprecipitated from cultures grown until early exponential phase using anti-FLAG antibodies followed by elution of the recombinant Rho-RNA complexes using a 3xFLAG peptide. A control Co-IP was performed using the untagged Rho protein from a WT strain. Since the Rho-3xFLAG protein was successfully enriched in the elution fraction (Fig. S12), the co-immunoprecipitated RNAs were extracted for sequencing. By comparing the RNA reads from Rho-3xFLAG purification and the untagged Rho, we identified 328 RNAs enriched with Rho (at least two-fold enrichment with padj < 0.05) (Table S2). The majority of the enriched RNAs (152 out of 328) are antisense RNAs or overlapping genes, correlating with the major function of Rho in suppressing pervasive transcription. To identify putative direct targets of Rho, we cross-referenced our RNA-seq and RIP-seq datasets, yielding 121 RNAs significantly enriched in RIP-seq and whose expression was affected by *rho* deletion (Table S3). This list includes eight mRNAs known to be regulated by the SaeRS TCS, such as *saeP*, *hlb-1*, *sbi*, *lukGH*, *scn*, and *chp* (Fig. 5B). Among other candidates, is *ribH* from the riboflavin synthesis operon, which is under the control of the FMN riboswitch (Table S3). Among the downregulated genes in the Δ*rho* mutant strain, only six transcripts co-immunoprecipitated with Rho, such as *narT* (nitrate transporter) and the operon including *cydB*, *ktrA* and BSR30_RS05090 genes.

In summary, our RIP-Seq data correlates well with our differential transcriptomics with a high selectivity for RNAs repressed by Rho since the expression of the vast majority (115/121) of enriched RNAs was highly upregulated in the *rho* mutant.

### Rho-dependent termination of specific mRNA targets

To monitor Rho effects on target RNAs, we constructed a GFP reporter with sequences containing the regulatory region of two target genes under the control of a constitutive promoter in the pCN57 vector^59^ (Fig. 7A). All plasmids were transformed into the strains HG001 pMKX (wild-type strain containing an empty pMKX plasmid), HG001 *△rho* pMKX (*rho* deletion mutant containing an empty pMKX plasmid), and HG001 *△rho* pMKX::*rho* (*rho* deletion mutant strain complemented with the pMKX::*rho* plasmid expressing *rho* under the control of a xylose-inducible promoter)^25^. As previously published, we observed a leaky expression of the *rho* transcript in the complemented strain without promoter induction (Fig. S13)^25^. The strains were grown in BHI medium until exponential phase and cells were lysed to measure GFP expression. If Rho terminates transcription at the predicted locations of the constructs, elongation through the *gfp* gene should be affected. Conversely, Rho transcriptional readthrough should lead to increased elongation through the *gfp* gene, resulting in higher expression of the GFP protein.

**Figure 7.**
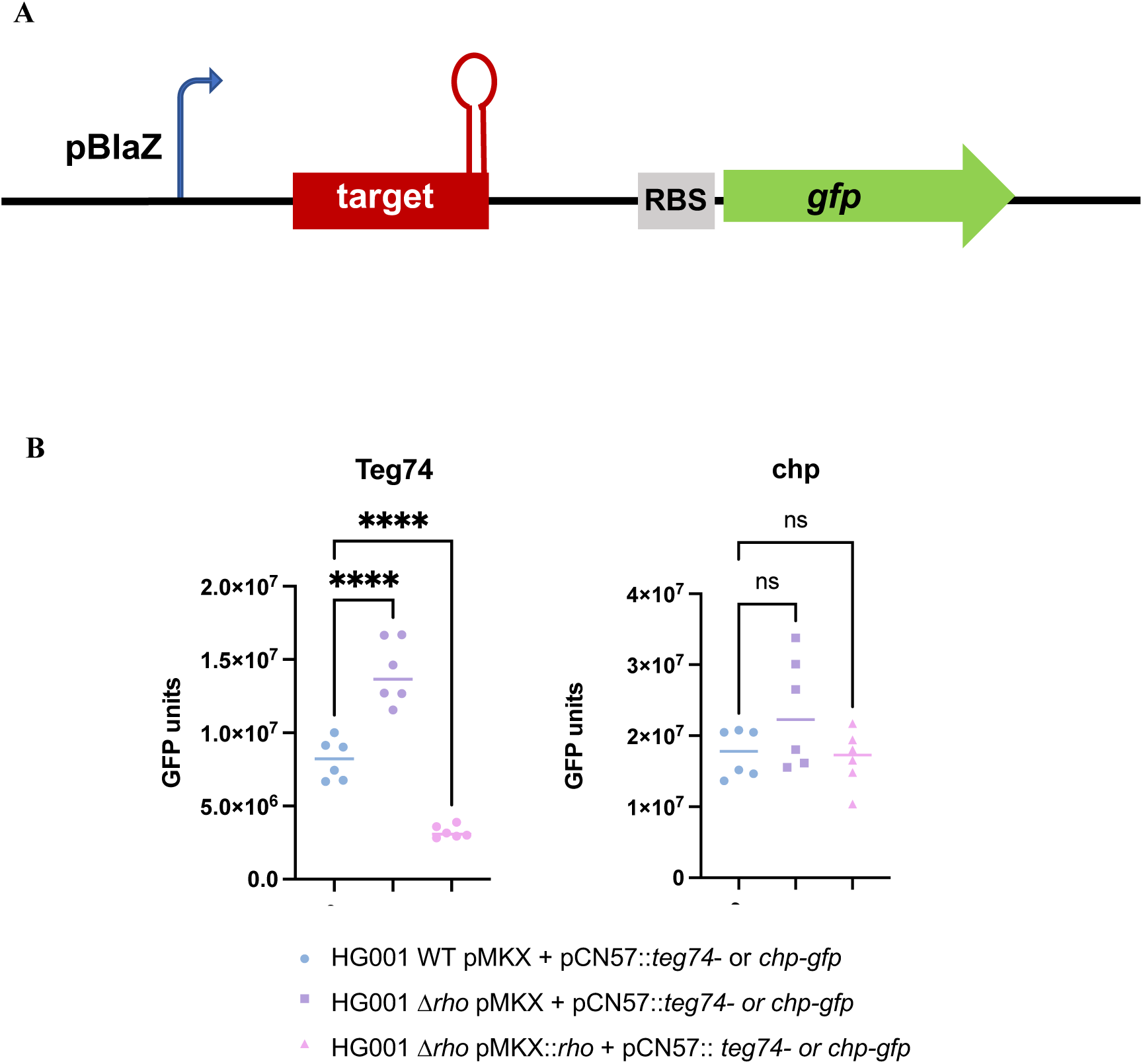
Rho terminates transcription of *teg74* and potentially *chp*. **A**. Schematic representation of pCN57::*gfp* reporter fusion plasmid. Target sequences of the *chp* and *teg74* genes comprising their own Rho-independent terminator hairpin are cloned upstream the *gfp* sequence under the control of the constitutive promoter BlaZ. RBS is for ribosome binding site. **B.** GFP levels of cellular extracts from strains HG001 WT pMKX, HG001*△rho* pMKX and HG001*△rho* pMKX::*rho* transformed with plasmids containing the sequences of *chp* and *teg74.* Cells were grown until exponential phase and lysed extracts were used for GFP measurement (n=6). Statistical analysis was performed with one-way ANOVA test.

To analyze the effect of Rho on the *teg74* riboswitch, we have first cloned the sequence of the riboswitch including the Rho-independent terminator in the reporter fusion construct. The data showed a significant increase of the GFP synthesis in the Δ*rho* mutant strain as compared to the WT and the complemented mutant strains (Fig. 7B). The same type of experiment was performed for the *chp* operon. In that case, we used a fragment comprising the 3’-end of the *chp* gene and the intergenic region downstream containing a Rho-independent terminator. We chose this region where transcriptional readthrough was observed (Fig. S9). The data revealed a tendency for a slight increase of GFP synthesis in the Δ*rho* mutant strain in comparison to the WT and complemented strains (Fig. 7B).

These data strongly support a direct role of Rho in transcription termination of specific mRNAs.

### Rho expression impacts *S. aureus* virulence

Based on the Rho-dependent regulation identified above, different phenotypic assays were chosen to define the impact of Rho on the physiopathology of *S. aureus*. We have compared the behaviors of the WT HG001 strain, the isogenic Δ*rho* mutant strain, and the complemented strain.

Since several mRNAs encoding exotoxins (Hla, HlgC) were upregulated in the Δ*rho* mutant strains, we first measured the amounts of the alpha (Hla) and gamma (HlgC) hemolysins by ELISA. A 2,8-fold increase in Hla and a 16-fold increase in HlgC were observed in the Δ*rho* mutant strain compared to the WT strain, and the mutant strain overexpressing Rho (Fig. 8A). These protein levels correlated with an increase in the cytotoxic activity of the Δ*rho* mutant strain on specific monocytes expressing gamma hemolysin receptors C5aR1 (3-fold increase) and CxCR2 (4-fold increase) (Fig. 8B) and on rabbit red blood cells sensitive to alpha hemolysin (4,6-fold increase) (Fig. 8C).

**Figure 8.**
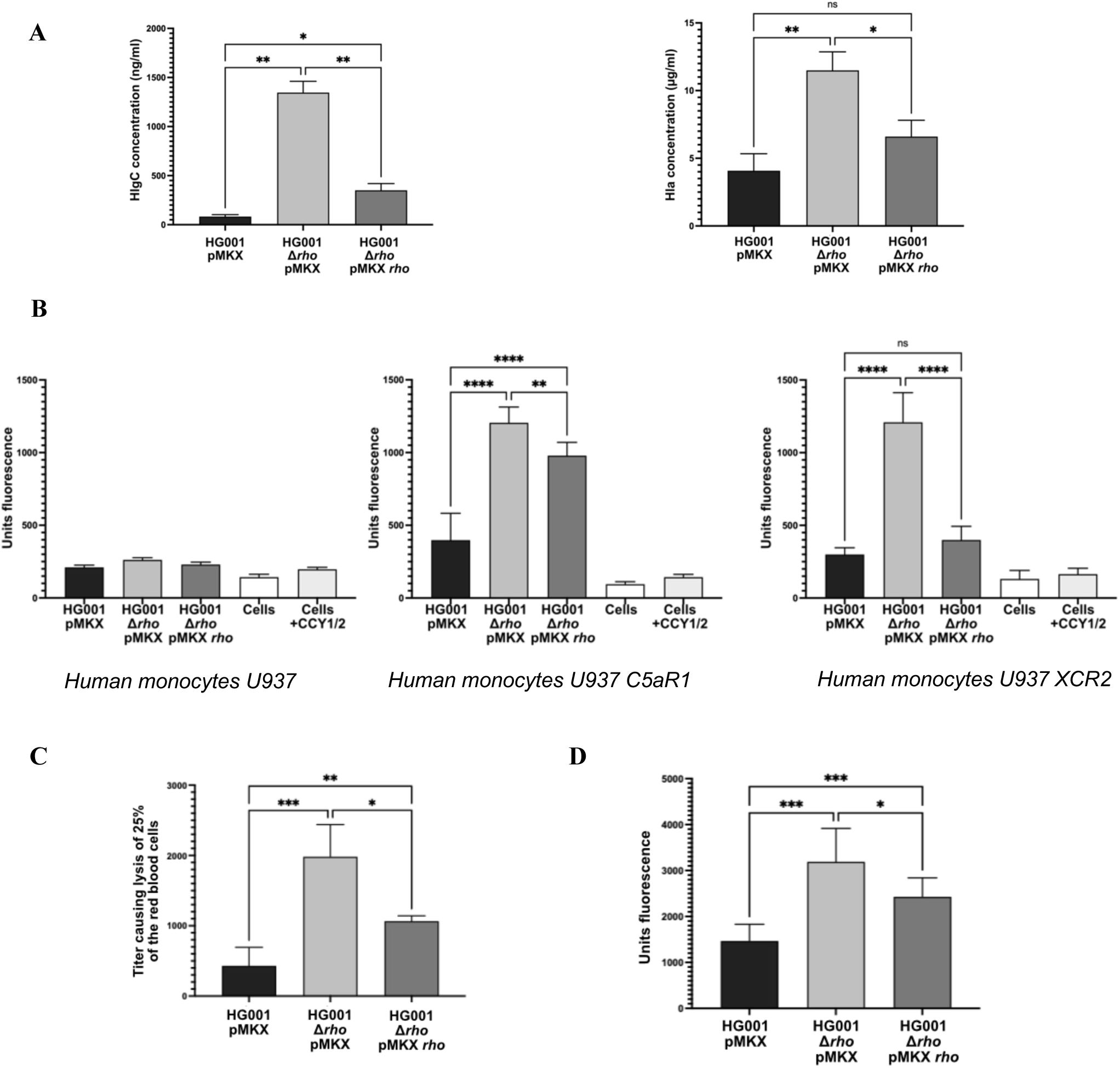
Impact of Rho on *S. aureus* cytotoxicity. **A.** Quantification of HlgC and Hla proteins (in nanograms per milliliter) from supernatants of HG001 WT (HG001-pMKX), mutant deleted for *rho* gene (HG001Δ*rho* pMKX) and complemented strain (HG001Δ*rho* pMKX::*rho*) cultures grown during 18 h in CCY medium (n=3). **B.** Bacterial supernatant cytotoxicity was quantified by measuring propidium iodide incorporation into U937 monocytes, and into U937 monocytes expressing gamma hemolysin receptors C5aR1, and CXCR2 (n=9). **C.** Graphic representation of the Hla hemolytic activity of the supernatants from cultures grown during 18 h in CCY medium on rabbit red blood cells (n=3); the Hla activity value corresponds to the inverse of the exact dilution causing the lysis of 25% of the red blood cells. **D.** Isogenic strains of *S. aureus* examined for their binding capacity to fibrinogen coated on 96-well plates, at concentration of 5 µg/ml. The adherence of bacterial cells was quantified at 485 nm using TECAN plate reader. Mann-Whitney tests performed: * = p-value < 0.05, ** = p-value < 0.01, *** = p-value < 0.001, **** = p-value < 0.0001.

Other mRNAs encoding several adhesin factors (Fibronectin binding proteins FnbAB, coagulase, and map) were also upregulated in the absence of Rho (Fig. 6). We first tested the fibrinogen adhesion phenotype *in vitro* and observed an increased ability of the Δ*rho* mutant strain to adhere to fibrinogen compared to the WT strain (Fig. 8D). Additionally, we monitored the ability of the same strains to coagulate rabbit plasma and detected a higher coagulase titer in the Δ*rho* mutant strain compared to the WT strain (Fig. S14). Noticeably, a restoration of most of the phenotypes was observed in the HG001Δ*rho*::pMKX-*rho* complemented strain, albeit with a reduced effect as compared to the WT strain. One cannot exclude that the over-expression of Rho from the pMKX plasmid resulted in a less active protein under the tested conditions.

All-in-one, these data show that Rho decreases cell cytotoxicity and reduces cell adherence reinforcing that Rho is an attenuator of virulence^26^.

## DISCUSSION

In this study, we investigated and characterized the regulation and the roles of the transcription termination factor Rho in *S. aureus*. This highly conserved protein is found in over 90% of bacterial genomes and accomplishes diverse biological functions, including regulation of specific operons, participation in sRNA-dependent mechanisms, and prevention of spurious transcription. In *E. coli*, Rho has a broad role, since it terminates transcription of up to 50% of all genes and often cooperates with NusA and NusG^21^. Rho can be part of the RNase E-based degradosome, demonstrating that Rho is a versatile and global regulator.^60^ In *S. aureus*, however, besides its role in controlling pervasive transcription, the functions of Rho appear to be more specific to virulence regulation. Indeed, the deletion or inactivation of Rho leads to increased expression of the SaeRS dependent-targets, resulting in enhanced virulence, as tested in a murine model of infection^26^. While virulence is a tightly controlled and time-dependent phenotype, we demonstrate that Rho synthesis is growth phase-dependent and is post-transcriptionally control by the quorum sensing regulatory RNAIII. In addition, we identify new Rho-dependent targets highlighting the dual functions of Rho as both a suppressor of pervasive transcription and a regulator of virulence gene expression.

### Rho protein synthesis is under the control of the quorum sensing dependent RNAIII

We showed that the expression of the *rho* mRNA levels decreases sharply at the entrance of the stationary phase. This regulation is under the control of the *agr* quorum-sensing system, specifically by its main effector, the 514-nt long RNAIII^37–41^. Indeed, the time-dependent regulation of the *rho* mRNA is alleviated in the deletion Δ*rnaIII* mutant strain. *In vitro*, RNAIII forms a stable complex with *rho* mRNA, which can be cleaved by RNase III (Fig. 3B, 4B). Our *in vitro* data support the existence of several interacting sites between RNAIII and *rho* mRNA, involving at least hairpin loops H7 and H9 of RNAIII (Fig. 3, S5C). While the deletion of either H9 or H7 only partially affected the binding to *rho*, their simultaneous deletion had a synergistic effect but did not completely abolish the binding. These data suggested that other compensatory binding sites between RNAIII and *rho* mRNA take place. Noteworthy, the 5’ UTR and the coding sequence of *rho* mRNA contain several redundant GGGA motifs potentially complementary to the UCCC motifs found in hairpin loops 7, 13 and 14 of RNAIII. This GGGA motif is also part of the binding sites of other RNAIII targets, such as *coa* and *rot* mRNAs, and a similar RNase III cleavage has been observed in the hairpin loop 7 at the C-rich motif due to a loop-loop interaction^39,41,61^. The importance of each hairpin of RNAIII for interaction with *rho* would need further clarification. We nevertheless propose that H9 of RNAIII binds to the coding region of *rho* (nts +46 to +82), forming at least 10 consecutive base-pairing interactions, further stabilized by hairpin loop H7 binding either to the _314_GGGAU_318_ or to the _327_UGGGA_331_ motif (Fig. S5C). We do not exclude that hairpin loops H13 and H14 of RNAIII may contribute to the overall stability of the complex by binding to the UGGG motifs in the 5’UTR of *rho* including its Shine and Dalgarno (SD) sequence. Since RNAIII covers a large region of *rho* mRNA next to the ribosome binding site, *rho* mRNA translation is expected to be repressed. Using toe-printing and *in vitro* translation assays, we showed that RNAIII binding to *rho* inhibits translation most likely by preventing the recruitment of the ribosome at the initiation step. We propose that the untranslated mRNA is more sensitive to degradation by ribonucleases. This would most probably contribute to the rapid depletion of *rho* mRNA at late exponential level of growth.

### Rho-dependent termination can enhance gene expression using different mechanisms

Deletion of *rho* increased the expression of antisense transcripts, whose functions are still controversial; consequently, the global effect of this increased pervasive transcription is difficult to evaluate. In our study, we did not always observe a correlation between an enhanced expression of an antisense and downregulation of the corresponding sense transcript. For instance, the antisense produced in the Δ*rho* mutant strain after readthrough at the *chp* gene did not have any effect on the levels of the sense transcript (Fig. S9 and Table S1A). Intriguingly, we often detected enhanced antisense transcription in poorly transcribed regions and we cannot exclude that the effects of antisense transcription are masked by the low basal levels of sense transcription. Nevertheless, the co-immunoprecipitation of many antisense transcripts with Rho suggests that this factor is needed to suppress their expression most likely through premature transcription termination and is in accordance with the roles of Rho in suppressing pervasive transcription.

Noteworthy, several genes involved in nitrate respiration were positively regulated by Rho. We frequently observed the generation of antisense transcripts to these genes, pointing to possible sense/antisense RNA regulation (Fig. S6). This hypothesis is supported by the fact that none of the relevant mRNAs co-immunoprecipitated with Rho in the RIP experiment, while several antisense transcripts to the affected genes were enriched (Table S1). Thus, Rho could inhibit the synthesis of these antisense transcripts to ensure the proper expression of genes implicated in nitrate respiration. However, we cannot exclude the possibility that the effect on these genes is indirect and occurs through other upstream pathways. In the RIP experiment, the enrichment of *narT* mRNA, which encodes a nitrate transporter, suggests a direct effect of Rho on this gene. In addition, a recent study indicates that the *narGHJI* operon is mostly expressed at the early exponential phase when *rho* mRNA is present^62^.

Besides nitrate respiration, Rho activates the synthesis of proteins (e.g., the dehydrogenases Adh and Ldh-1, Hmp, and the SrrAB TCS) involved in anaerobia although *rho* transcription does not rely on SrrAB known to sense low O_2_ levels (Fig. S1). All these genes have in common that they are transcriptionally controlled by the redox sensing regulator Rex^63^. When O_2_ level is high, Rex is activated by NAD^+^ and represses anaerobic genes such as dehydrogenases. Nonetheless, *rex* transcript is not differentially expressed in the Δ*rho* mutant nor pulled out in the RIP-seq experiment with Flagged-Rho. We can postulate that for a yet unknown reason, the NADH/NAD^+^ ratio is perturbed in the Δ*rho* mutant which would interfere with Rex activity. The expression of anaerobic gene enables *S. aureus* to adapt to changes in oxygen concentration, which is especially relevant in an infection context. Since oxygen levels can vary within the human host, these variations can impact bacterial physiology and virulence factor production^64^. Therefore, regulation of anaerobic gene expression by Rho could influence the ability of *S. aureus* to adapt to different host tissues and its pathogenesis, particularly in low oxygen environments such as abscesses. Similarly to *narT*, the *cydB*, *ktrA* and BSR30_RS05090 mRNA operon involved in respiration and potassium transport were also co-immunoprecipitated with Rho. These data could indicate that for a reduced number of genes (mostly involved in metabolism/anaerobic respiration), Rho might be required for proper transcription termination.

### Rho contributes to FMN-responsive transcriptional riboswitch regulation

In bacteria, homeostasis of riboflavin or vitamin B2 is regulated by the widely distributed class of flavine mononucleotide (FMN) riboswitches. The FMN riboswitches regulate the biosynthesis and uptake pathways of riboflavin, which is then converted into (FMN) and flavine adenine dinucleotide (FAD), two essential co-factors for bacterial growth as they are critical for many redox reactions of cellular metabolism^65^. FMN acts as the primary ligand regulating FMN riboswitches by binding to the aptamer when its concentration is high. Binding induces conformational changes to inhibit either the transcription or the translation of the downstream genes^66^. In *S. aureus*, inhibiting riboflavin biosynthesis does not reduce virulence, but simultaneously disrupting biosynthesis and uptake pathways with FMN riboswitch inhibitors like roseoflavin or ribocil-C efficiently impairs MRSA growth and virulence in a murine septicemia model^67^. Thus, specifically targeting FMN riboswitches in *S. aureus* could be a promising strategy to treat drug-resistant bacteria.

*S. aureus* encodes two highly related but different FMN-responsive riboswitches. Teg74 regulates the riboflavin biosynthetic operon (*ribDEBAH*) at the transcriptional level while Teg66 controls the uptake transporter (*ribU*) at the translational level. Only, the FMN-responsive transcriptional riboswitch *teg74* gene along with the downstream *rib* operon were upregulated in the Δ*rho* strain. The transcripts were also enriched upon RIP, although with a non-significant padj value, which may be explained by the relatively low expression of the operon in rich growth medium and early exponential phase. Using the reporter gene where *teg74* was cloned upstream the *gfp*-encoding sequence, a significant increase of GFP synthesis was observed in the Δ*rho* mutant strain. These observations strongly suggest that Rho-dependent termination of *teg74* is required for the repression of the downstream *rib* operon, linking Rho-dependent termination to riboswitch-mediated regulation in *S. aureus*. Although both riboswitches share 96% identity^67^, Teg66 and *ribU* were not affected upon *rho* deletion nor co-immunoprecipitated with Rho, suggesting the specific action of Rho on the transcriptional FMN riboswitch. Interestingly, the association of Rho to this particular riboswitch is conserved in different bacteria such as *E. coli*^68^, *Clostridium botulinum*^69^ and *Bacillus subtilis* (E. Bidnenko, S. Durand, personal communication).

Overall, Rho-dependent regulation of the transcriptional FMN riboswitch in *S. aureus* might contribute to modulate pathogenesis.

### Rho acts as an attenuator of virulence

The RNA-seq analysis together with the RIP-seq revealed that the absence of Rho lead to an increased production of virulence factors, explaining the phenotypic assays (Fig. 8). We mainly focused on virulence associated genes and only considered the direct targets of Rho that are repressed. Since Rho attenuates virulence in *S. aureus*, our rationale is that Rho induces premature transcription termination of these targets to reduce their expression. Since almost the entire *sae* regulon is upregulated in the absence of Rho, it was stipulated that the effect of Rho would occur through the two-component system SaeRS, which acts as a major regulator of virulence gene expression in staphylococci^26^. Our data indicates that Rho binds only to the *saeP* mRNA, one component of the *sae* operon. The *saePQRS* operon contains two promoters: P1 induces the expression of *saeP* and P3 the expression of *saeR*, and *saeS*. The P1 promoter is strongly autoregulated, and is activated by the quorum sensing effector AgrA, while it is repressed by Sigma factor B^70^. The *saeP* transcript can be produced by two independent mechanisms: one involves a specific RNase Y cleavage that generates an unstable *saeP* and a stable *saeQRS* mRNAs, and the other involves a termination event at a Rho-independent terminator that produces *saeP*^71^. Our data suggested that in the Δ*rho* mutant, a transcriptional readthrough event occurred at *saeP* allowing further transcription into the *saeQRS* genes (Fig. S7). Thus, *saeP* can be transcribed separately from *saeQRS* as previously hypothesized^71^. The fact that only *saeP* mRNA co-immunoprecipitated with Rho supports the idea that Rho might terminate the transcription of *saeP*. Hence, it is possible that premature termination and/or degradation of *saeP* alone could significantly impact the functioning of this system. Interestingly, in *S. aureus*, PNPase negatively alters the expression of the *saePQRS* operon^72^ in the absence of maturation by RNase Y that forms protective 3’ extremities^72^. Thus, it is tempting to propose that Rho-dependent termination of *saeP* might induce degradation by PNPase affecting expression of the entire operon as observed in the differential transcriptome analyses (Table S1). Meanwhile, SaeRS autoinduces the transcription of the entire *sae* operon at P1 promoter. This feedback loop likely generates sufficient SaeRS TCS in the activated state. We propose that the Rho-dependent repression of *saeP* might play a critical role in disrupting this feedback loop, thereby significantly reducing the expression levels of the entire *sae* operon (Fig. 9). Intriguingly, the deletion of *saeP* or the P1 promoter has been shown to enhance *S. aureus* cytotoxicity against human neutrophils by modulating the *sae* regulon^73,74^. Consequently, the deletion of either *saeP* or *rho* generates similar phenotypic effects due to the activation of the *sae* regulon. SaeP has been demonstrated to form a complex with SaeQ *in vitro*, which activates the phosphatase activity of the sensor kinase SaeS^75^. This could explain the seemingly indirect effects observed for some *sae*-regulated genes, such as *sbi*, where a major transcriptional effect was detected without any visible readthrough event (Fig. S8). Thus, overexpression of SaeRS targets in the Δ*rho* mutant, especially of those that were not enriched upon RIP, might be indirect through *saeP*. Most importantly, *rho* expression must be repressed in the stationary phase when SaeRS is active (see below and Fig. 9).

**Figure 9.**
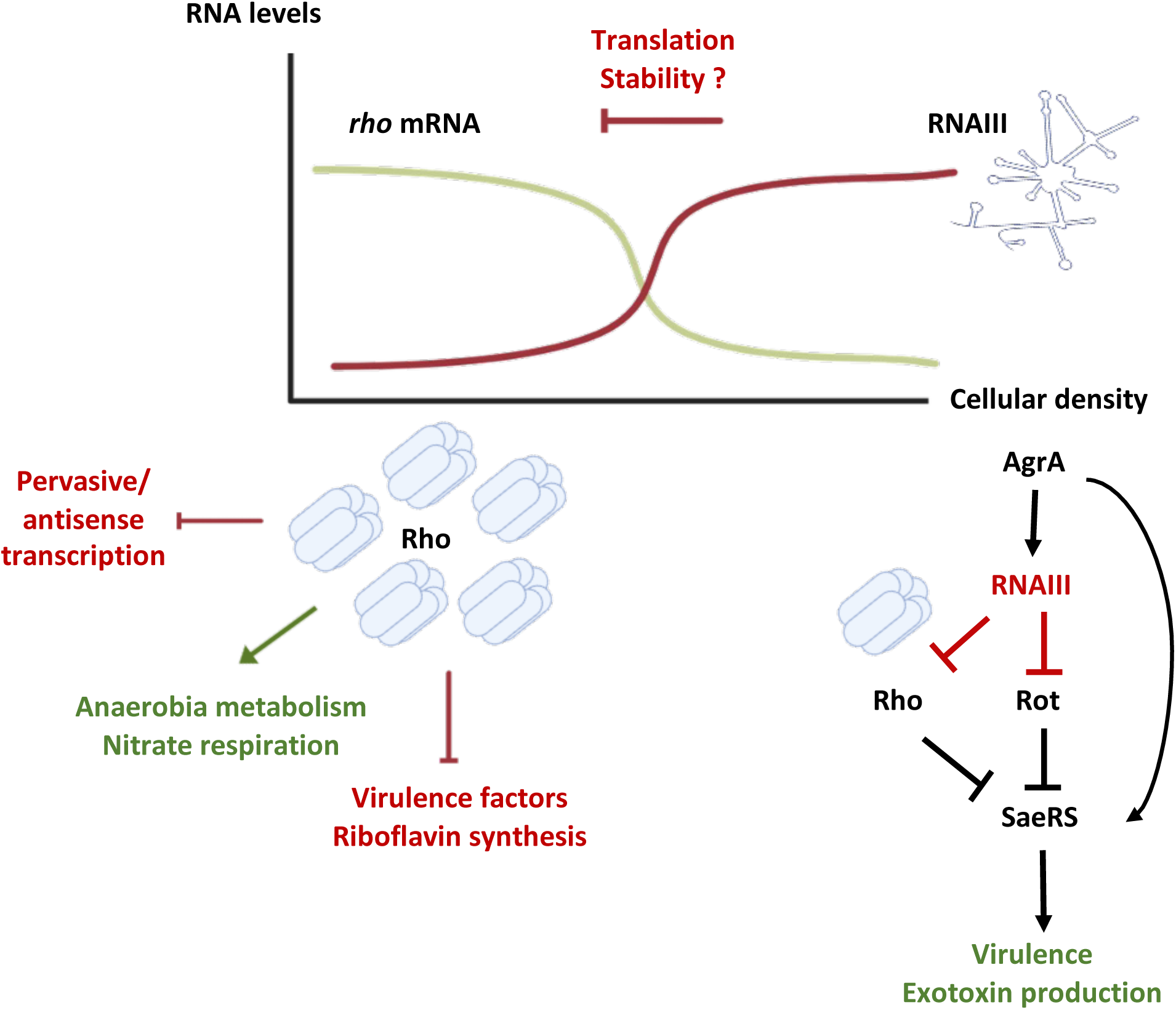
Proposed model of regulation of *rho* during *Staphylcoccus aureus* growth. At low cell density, Rho play crucial roles by repressing the expression of the *sae* operon, suppressing pervasive antisense transcription, and activating the synthesis of proteins involved in anaerobic and nitrate metabolisms. At high cell density, the quorum sensing induced RNAIII represses the translation of *rho* mRNA through direct RNA-RNA pairing. The expression of *sae* operon is enhanced directly via AgrA, and indirectly via RNAIII. These regulatory mechanisms collectively promote the synthesis of exotoxins. Activation is represented by arrows and repression by bars: red color is for post-transcriptional regulation and black color for transcriptional control. Green arrow depicts the direct and indirect effect of Rho on anaerobia and nitrate respiration (see text for details).

However, several of the SaeRS targets also co-immunoprecipitated with Rho pointing to a simultaneous indirect and direct effects of Rho (Fig. 6B). The genomic region including *chp* has retained our attention, because a readthrough was observed on both strands downstream the *chp* and BSR30_RS10675 genes, and because many of the depicted genes were enriched in the RIP-seq (Fig. 6 and Table S1). We believe that Rho is involved in premature transcription termination of several loci in this region, for instance at the *chp*, BSR30_RS10675 and *scn* genes, containing Rho-independent terminators and also on the *hlb_1* gene with no predicted intrinsic terminator hairpin. Using a transcriptional GFP reporter plasmid fused to the *chp* gene, we identify a readthrough event due to the absence of Rho *in vivo*, supporting the direct involvement of Rho in the regulation of the *chp* gene. All these genes are virulence factors and an increase in their mRNA levels may explain the hypervirulence phenotype of the deletion mutant of *rho*^26^.

Considering the role of Rho as an attenuator of virulence, the repression mediated by quorum sensing-induced RNAIII at high cell density represents an important asset for pathogenicity. Besides RNAIII represses the translation of both *rho* mRNA and *rot* mRNA in a very similar manner^51^, the latter encoding a transcriptional repressor of SaeRS^76^. Additionally, the transcription of the *sae* operon was shown to be enhanced by AgrA. Taken together, these regulatory mechanisms contribute to optimize the synthesis of exotoxins facilitating the invasive state of the bacterial population (Fig. 9). Moreover, a recent work showed that the RNAIII-mediated translational repression of Rot is essential to protect the bacteria from exogenous reactive oxygen species^77^. Interestingly, this long-lived protection was uncoupled from quorum sensing activation suggesting that the bacteria anticipate the ROS-mediated immune attack and the oxidative damages. As Rho protein regulates genes involved in riboflavin synthesis, nitrate respiration and anaerobia, it is tempting to propose that the RNAIII-dependent repression of *rho* might also be required under specific conditions encountered during the infection, even if RNAIII is not highly expressed. Indeed, it was previously shown that efficient repression of target genes occurred as soon as RNAIII is induced^78^.

### Rho protein is involved in phenotypic switches in different bacterial species

Interestingly the growth phase dependent repression of Rho is also observed in *B. subtilis,* and is required for cell adaptation, although the mechanism of regulation is not known^24,79^. In *Acinetobacter baumanii* which can adopt a virulent and an avirulent phenotypes, it has been shown that Rho is necessary for premature termination of transcriptional regulators and their repression in the virulent status^33^. Conversely, when switching to an avirulent phenotype, Rho-dependent termination is reduced to allow correct expression of these regulators. Interestingly, levels of Rho remained unchanged in both phenotypes, suggesting the existence of mechanisms that inactivate the protein or reduce its binding to the mRNA targets^33^. As demonstrated in *E. coli,* Rho can be transiently and reversibly inactivated through oligomerization and aggregation^32^. Also, in *Clostridium botulinum,* Rho can adopt free or aggregated conformations due to the presence of a prion-like domain in the NTD which can impact the activity of the protein and act as a regulatory switch of Rho functions^80^. Further analyses will be needed to investigate if a similar phenomenon exists in *S. aureus,* since Rho also contains an NTD insertion that could mediate a similar regulatory switch. This might be relevant because the levels of the Rho protein slightly vary between exponential and stationary phase, suggesting that other mechanisms might exist to ensure the inactivation of Rho at high cell density. Nevertheless, the RNAIII-dependent regulation of *rho* is expected to enhance the synthesis of several exotoxins at this stage. This is also supported by the phenotypic assays showing that Rho decreases cell cytotoxicity.

## CONCLUSION

The role of Rho in the control of genome surveillance through the regulation of antisense transcription has been conserved among bacteria. These pervasive transcripts are often associated to horizontally acquired genetic material or mobile elements. In *B. subtilis* and *S. aureus* accessory genes have been acquired for sporulation or virulence, respectively, which would explain why Rho seems to have more specialized regulatory functions in these bacteria. This ubiquitous termination factor has evolved for the adaptation of Bacillota to the physiological remodeling taking place in the transition to stationary phase. Our findings have also unveiled the *rho* mRNA as a novel target of RNAIII, adding another level to the complex networks controlling the activation of exotoxins during stationary phase. Overall, our results give a first glimpse of the roles and functions of Rho in *S. aureus* and placed Rho as another key component of the main regulatory networks of virulence of *S. aureus*.

## MATERIALS AND METHODS

### Strains and plasmids

All strains and plasmids used and constructed in this study are described in Tables S4 and S5. The primers designed for cloning and mutagenesis are provided in Table S6. *E. coli* strain IM08B was used as an intermediate strain for plasmid amplification before electroporation in *S. aureus*. Plasmids were extracted from chemically transformed *E. coli* pellets with the NucleoSpin Plasmid kit according to the manufacturer’s protocol (Macherey-Nagel). Electrocompetent *S. aureus* strains were then transformed with the purified plasmids by electroporation (Bio-Rad Gene Pulser). The HG001 *rho* and HG001 *rnaIII* deletion mutants as well as the HG001 *rho-3xflag* strain were constructed by homologous recombination using plasmid pMAD in *S. aureus.* HG001Primers used are listed in Table S6.

The pCN51-P3::*rnaIII* and P3::MS2-RNAIII plasmids were constructed by ligating PCR-amplified fragments (see Table S6 for primers) containing the 514 nt-long sequence of *rnaIII* previously digested with *Sph*I/*Bam*HI or the sequence of *rnaIII* fused to an MS2 tag at the 5’end digested with *Pst*I/*Bam*HI into a pCN51-P3 plasmid digested with the same enzymes, respectively. The pCN51-P3::*rnaIII△H7*, pCN51-P3::*rnaIII△H9* and pCN51-P3::*rnaIII△H7△H9* plasmids containing deletion mutants of hairpins 7, 9 or both, of RNAIII were constructed by site directed mutagenesis using plasmid pCN51-P3::*rnaIII* as a template. A PCR using Phusion High Fidelity DNA polymerase (Thermo Scientific) was performed using the primers listed in S6 to introduce the deletions. One denaturation step (30 s at 90°C) was followed by eighteen cycles of amplification of the fragment (30 s at 95°C, 1 min 55°C, 6 min at 68°C) and ultimately an elongation step (5 min at 72°C) was performed. PCR products were then treated with 10U/µL *Dpn*I to eliminate the template pCN51-P3::*rnaIII* plasmid. Reactions were then transformed into *E. coli* TOP10 and positive clones were selected and sequenced. These plasmids were used as templates for PCR amplification to introduce the T7 promoter for templates in *in vitro* transcription assay. Plasmids pCN57::GFP-*chp,* pCN57::GFP-*teg74,* pCN57::GFP-*saeP* and pCN57::GFP-*teg66* were constructed by ligating PCR-amplified fragments of the *chp*, *teg74, saeP and teg66* genes (see Table S6 for primers) previously digested with *Eco*RI/*Bam*HI into a pCN57::GFP plasmid digested with the same enzymes.

### Growth conditions

*E. coli* strains were grown in Lysogeny-Broth (LB, Roth) medium supplemented with ampicillin (100 μg/mL). *S. aureus* strains were grown in Brain-Heart infusion (BHI, BD) media containing 10 μg/mL of erythromycin or chloramphenicol when needed.

### pMAD mutagenesis

Strains HG001 *△rho*, HG001 *△rnaIII* and HG001 *rho-3xflag* were constructed by allelic recombination using the shuttle vector pMAD as previously described^81^. To generate the mutants, we amplified fragments of the flanking upstream and downstream regions of the targeted genes by PCR using chromosomal DNA of *S. aureus* HG001 as template. The final PCR products were obtained by flanking PCRs or double digestion/ligation steps (Table S6 for primers) and cloned into pMAD shuttle plasmid with *Bam*HI/*Bgl*II or *Bam*HI/*Eco*RI restriction enzymes. Plasmids were amplified into RN4220 after electroporation and then transferred in HG001 for homologous recombination steps as described in Arnaud *et al*, 2004^81^.

### Northern blot

Isolation of total RNA of *S. aureus* cultures was performed accordingly to the FastRNA pro blue kit (MP Biomedicals) using the FastPrep machine (MP Biomedicals) for mechanical lysis of bacterial cells. Electrophoresis of 15 µg of total RNA was run in 1% agarose-TBE 1X gel containing 25 mM guanidium thiocyanate (Sigma). After migration at 150 V, RNAs were transferred on Hybond+ nitrocellulose membrane (Cytiva) by capillarity with saline-sodium citrate 2X buffer (SSC 2X) and then fixed by UV-crosslinking (Stratalinker 1800 Stratagene). RNA detection was performed by hybridization with specific digoxigenin (DIG)-labeled RNA probes complementary to each targeted RNA as described in Tomasini *et al*, 2017^46^. (see S3 for primers used for DIG-probe production).

### Western blot

Total proteins were extracted from *S. aureus* cultures grown for 2, 4 or 6 h in BHI. After harvesting by centrifugation, bacterial pellets were washed once wit 1X PBS, then resuspended in 1X PBS and finally lysed using the Fastprep (MP Biomedicals). Quantification of proteins was performed with Bradford reagent following the manufacturer protocol (BioRad) and using BSA as standard. 10 µg of total protein extract were migrated on 12 % polyacrylamide-SDS gel and transferred onto a PVDF membrane using the Trans-blot Turbo Transfer system (BioRad). Membranes were incubated with anti-FLAG mouse antibodies (Sigma) at the final dilution of 1:2500 followed by the incubation with a goat anti-mouse peroxidase (HRP) antibody (BioRad) at the final dilution of 1:5000. Membranes were revealed using the ChemiDoc (BioRad) and were then stained by Coomassie blue as loading controls of samples.

### MAPS

Cultures of HG001 pCN51::MS2 or *△rnaIII* pCN51::MS2-RNAIII strains were grown in BHI medium for 6 h and then centrifuged. Pellets were then harvested, lysed and purified by affinity chromatography as previously described^82^. RNA was isolated, purified from the collected fractions with PCI (Carl Roth, pH 4,5-5), precipitated with EtOH and used for Northern blot^82^.

### Transcriptomics

Cultures of HG001 and HG001*△rho* strains were grown in BHI medium until OD_600_=0.5 and harvested by centrifugation. Total RNA was extracted as described above and treated with DNase I (Sigma) prior to RNA-seq. RNAs were treated to deplete abundant rRNAs and the cDNA libraries were performed using the NEBNext Ultra II directional RNA kit. The libraries were sequenced using NextSeq2000 P2. Each RNAseq reaction was performed in duplicates. The reads were then processed to remove adapter sequences and poor-quality reads using Trimmomatric^83^, then converted to FASTQ format using FASTQ Groomer^84^, and aligned to the HG001 genome^85^ using Bowtie2^86^. Finally, the number of reads mapping to each annotated feature was determined with HTSeq^87^. To perform the differential expression analysis, we used DESeq2^88^. All processing steps were performed on the Galaxy platform^89^.

### RIPseq

Cultures of HG001 WT and HG001 *rho-3xflag* strains were grown in BHI medium until early exponential phase (OD_600_=0,5) and harvested by centrifugation. Pellets were resuspended in lysis buffer (50 mM Tris-HCl pH 8, 150 mM NaCl, 1 mM EDTA, 1% Triton X-100 and protease inhibitors) and transferred into flacon tubes containing Lysing Matrix B silica beads (MP Biomedicals) to proceed to lysis of the cells using the Fastprep (MP Biomedicals). After centrifugation to remove cell debris, the supernatant was incubated for 2 h at 4°C on a rotator with pre-equilibrated ANTI-FLAG M2 affinity Gel (Sigma) as described by the manufacturer protocol. The suspension was centrifuged, the supernatant discarded, and the beads were washed twice with cold 1X TBS. The antibody-bound fusion protein Rho-3xFLAG was then eluted by competition with 100 µg/mL of triple FLAG peptide (Sigma) in 1X TBS. A sample of each fraction was used for Western blot to assess the enrichment of the tagged protein after elution. RNA was purified from the elution fraction with PCI precipitation (Carl Roth, pH 4.5-5) and treated with DNase I (Sigma) prior to RNA-seq analysis. RNAs were treated to deplete abundant rRNAs and the cDNAs were performed using the NEB Next Ultra II directional RNA kit. The libraries were sequenced using HiSeq 1000 Single read 50 bases. Each RNAseq was performed in duplicates. Treatment of RNAseq data was performed as described above. To identify co-precipitating asRNAs, the RNAseq analysis was analyzed as described above but in a reverse orientation.

### Preparation of RNAs for *in vitro* experiments

Transcription of the different RNAs used (RNAIII, RNAIII△H7, RNAIII△H9, RNAIII △H7△H9, *rho5’* and *rho3’*) was achieved using PCR products containing the sequence of the RNA downstream the T7 promoter, directly introduced with the primer (Table S6). These PCR products were used as templates for *in vitro* transcription using T7 RNA polymerase. In the case of RNAIII transcription, reactions were supplemented with 5 mM of ApG RNAs to avoid the 5’-end dephosphorylation step. RNAs were treated with DNase I and purified on an 8% polyacrylamide-8 M urea gel and eluted with 0,5 M ammonium acetate pH 6.1, 1 mM EDTA, and 0.1% SDS and precipitated in cold absolute ethanol. Precipitated RNAs where then labeled with T4 polynucleotide kinase (Fermentas) and [γ^32^P] ATP and ultimately purified on a 8% polyacrylamide-8M urea gel and eluted as described above.

### EMSA

The 5’-end radiolabeled RNAs (10’000 cps/sample, <1pM), and cold RNAs were denatured separately by incubating at 90°C for 1 min in 100 mM Tris-HCl pH 7.5, 300 mM KCl, 200 mM NH_4_Cl, then cooled down for 1 min on ice and renatured at 20°C for 10 min after addition of 10 mM MgCl_2_. Complexes were formed at 37°C for 15 min. After the addition of 1 volume of glycerol blue, the samples were loaded on a native 6% polyacrylamide gel containing 10 mM MgCl_2_ and migrated at 300 V and 4°C in 1X TB buffer with 10 mM MgCl_2._

### RNase III cleavage

The 5’-end radiolabeled RNAIII (10’000 cps/sample, <1 pM) and cold *rho* mRNA were denatured separately at 90°C for 1 min, cooled down on ice for 1 min and renatured in 1X TMK buffer (Tris HCl 100 mM pH7, MgCl_2_ 100 mM, KCl 1M) at 20°C for 10 min. Cold RNAs are added to the labeled-RNA mix and RNA-RNA complexes are formed at 37°C for 15 min. RNAs are then incubated with increasing concentrations of purified *S. aureus* RNase III (25 nM, 50 nM, 100 nM and 200 nM) in presence of Mg^2+^ or Ca^2+^ and 1 µg of yeast total tRNA at 37°C for 15 min. RNAs are purified with PCI precipitation (Carl Roth, pH 4.5-5) and loaded on a 12 % polyacrylamide-8 M urea gel.

For the alkaline ladder, radiolabeled RNA is incubated with Alkaline hydrolysis buffer 1X (Ambion) at 90°C for 10 min before addition of 1 volume of loading buffer (Ambion). For the T1 ladder, radiolabeled RNA is denatured at 90°C for 1 min in sequencing buffer 1X (Ambion). After renaturation at 20°C for 1 min, 1 U of RNase T1 (Sigma) is added and incubated at 37°C for 5 min before addition of one volume of loading buffer (Ambion).

For RNase III cleavage experiments with RNAIII mutants, unlabeled *rho* and RNAIII variants to final concentrations of 400 nM and 200 nM, respectively, were used to form duplexes and RNase III cleavage was performed as described above. Cleavage signals on RNAIII were detected by primer extension. Precipitated RNAs were then hybridized with a 5’ end-labeled primer complementary to the RNAIII 3’ end (Table S6) and reverse transcription with AMV reverse transcriptase was performed at 37°C for 30 min. Sequencing reactions were performed using DNA template for each tested RNA and Vent (exo)-DNA polymerase (NEB) (25 cycles, 1 min denaturation at 95°C, 1 min annealing at 52°C and 1 min elongation at 72°C).

### Toe-printing

The preparation of *S. aureus* 30S subunits, the formation of a simplified translational initiation complex with mRNA, and the extension inhibition conditions were performed as previously described^44^. Increasing concentrations of either RNAIII or RNAIIIΔ7Δ9 concentrations were used to monitor their effects on the formation of the initiation complex with *rho*.

### *In vitro* translation assays

The translation assays were performed using the 5’-*rho* mRNA and the PURExpress In vitro Protein Synthesis kit (NEB) supplemented with 70S staphylococcal ribosomes purified as described^90^. The construct carries an additional sequence corresponding to the FLAG peptide, which was inserted at the 3′ end of *rho*. The reactions were performed at 37°C for 2 h following the kit protocol. Experiments were also carried out in the presence of increasing concentrations of RNAIII or of RNAIIIΔ7Δ9. Reactions were stopped by adding an equal volume of Laemmli buffer. The proteins were detected by western blot with an antibody against the FLAG tag (Sigma).

### GFP fusions

*S. aureus* strains containing the pCN57 plasmids derivatives were grown in BHI medium supplemented with 10 μg/mL erythromycin and chloramphenicol until exponential phase. Total proteins were extracted and quantified as described above. For each sample, 200 µL of dilutions of 1 mg/mL, 0.5 mg/mL and 0.25 mg/mL of total proteins were loaded on a black uCLEAR 96 wells-plate (GreinerBioOne) and GFP signal was measured with the platereader Spectramax Id3.

### HlgC and Hla quantification by ELISA

Bacterial strains were cultured in CCY broth supplemented with chloramphenicol at 10 μg/mL at 37°C overnight with shaking to stimulate toxins production^91^. Samples were centrifuged at 10’000 g for 10 min and supernatants were collected. HlgC and Hla quantification was performed by a sandwich ELISA using custom-made antibodies. 96-well Nunc MaxiSorp plate (Thermo Scientific) were coated with anti-HlgC monoclanal (R&D Biotech) or anti-Hla polyclonal (MedImmune) antibodies at 10 μg/mL and 2 μg/mL respectively, and incubated overnight at 20°C. After 5 consecutive washes with phosphate-buffered saline (PBS)–0.05% Tween (PBS-T), wells were saturated for 1 h 30 min at 20°C with a blocking solution containing PBS-T, low-fat milk (5 g/L), and bovine serum albumin (BSA) (1 g/L). Standard dilutions from 15 to 1’000 ng/mL of recombinant HlgC or from 0,3125 to 20 ng/mL of recombinant Hla or the culture supernatant were denatured for 1 h at 95°C, loaded in duplicate, and incubated for 2 h at 37°C. For HlgC quantification, after washing, polyclonal rabbit F(ab)′_2_ biotinylated antibody (1.55 μg/mL, R&D Biotech) was added to each well, and the plate was incubated for 1 h 30 at 37°C and washed. Then, ExtrAvidin-peroxidase antibody (Sigma) targeting the biotin molecule and conjugated with horseradish peroxidase (HRP) was added. The plate was incubated for 1 h at 20°C.

For Hla quantification, the plate was washed, monoclonal anti-Hla antibody (2 μg/mL) was added, and the plate was incubated for 1 h 15 at 37°C. After washing, polyclonal goat F(ab)′_2_ ant-human Fcg-HRP conjugated with horseradish peroxidase (HRP) antibody (Jackson Immuno-Research) was added and the plate was incubated for 1 h 15 min at 37°C. For both, a final wash was performed and 75 μL of the substrate tetramethylbenzidine (KPL SureBlue; SeraCare) were added. The reaction was stopped with 1 N sulfuric acid. The plates were read at 450 nm in a Bio-Rad model 680 microplate reader.

### Red blood cells lysis

Bacterial strains were cultured in CCY broth supplemented with chloramphenicol at 10 μg/mL at 37°C overnight with shaking, centrifuged at 10’000 g for 10 min and supernatant were collected. Activity of alpha-hemolysin was confirmed by cell permeability assays using rabbit erythrocytes. In 96-well round bottom plate, bacteria culture supernatants were diluted in PBS (1/2 to 1/2048) and mixed with 50 µl of washed rabbit erythrocytes solution (10^8^ cells/ml; Atlantis). After 60 min of incubation at 37°C, the plate was centrifuged and the OD_415nm_ of the supernatants measured. The cut off (corresponding to lysis of 25% of red blood cells) was determined using the following formula: (OD lysis control – OD neg control)/2, where the lysis control is the total lysis of red blood cells with pure water and the negative control is the mix of red blood cells with PBS. For each condition, the HLA activity value corresponds to the inverse of the exact dilution causing the lysis of 25% of the red blood cells.

### Cytotoxicity assay

Bacterial strains were cultured in CCY broth supplemented with chloramphenicol at 10 μg/mL at 37°C overnight with shaking, centrifuged at 10’000 g for 10 min and supernatant were collected. U937 Human monocytes expressing C5aR1 or CXCR2 receptor^92^ were routinely cultured in Roswell Park Memorial Institute (RPMI ; Thermofisher) 1640 Medium supplemented with 10% foetal bovine serum at 37°C with 5% CO_2_. U937 cells were routinely cultured in DMEM growth medium supplemented with 10% foetal bovine serum at 37°C with 5% CO_2_. Cells were diluted at 1.10^6^ cell/ml and Iodure Propidium (IP) was added for a final concentration at 25 µg/mL. A 96-well plate was seeded with 90 µl of this solution and 10 µl of culture supernatant diluted at ½ in DMEM medium were added. IP incorporation into cells was measured by determining OD at 635 nm after excitation at 535 nm using TECAN plate reader. The measure was done every 10 min for 24 h and the OD max was conserved to compare cytotoxic activities of the different bacterial supernatants.

### Adherence assay

Adherence of bacterial cells to immobilized fibrinogen was performed as described by Hartford *et al*,1997^93^. Briefly, 96-well plate was coated with 5 μg/mL of fibrinogen (SIGMA) in PBS and incubated overnight at 4 °C with shaking. Bacterial strains were cultured in BHI broth at 37°C overnight. The plate was incubated with bovine serum albumin solution (1% in PBS) for 1 h at 37 °C. 1 mL of bacterial cell adjusted at 10^8^ CFU/mL were labelled with 2 µL of BacLight RedoxSensor Green Vitality (Thermofischer) for 15 min at 37°C with gentle shaking. 100 µL bacterial cell suspension was added per well in duplicate and the plate was incubated for 1 h at 37°C in the dark. After incubation, the plate was washed three times with PBS and the adherence of bacterial cells was quantified at 485 nm using TECAN plate reader.

### Coagulase quantification

Bacterial strains were cultured 5 h at 37°C with agitation in Brain Heart Infusion (BHI; Difco) supplemented with chloramphenicol at 10 μg/mL and then centrifuged. Coagulase activity was assayed by adding 0.5 mL of two-fold serial dilutions of culture supernatant in PBS to 0.5 mL of rabbit plasma (Biorad) diluted 1 in 3 in distilled water. The titer was the reciprocal of the highest dilution of the supernatant that showed evidence of clotting after incubation for 18 h at 37°C.

### Statistical Analysis

The statistical analyses were performed using GraphPad Prism 10 software. Data from 3 strains were compared using multiple comparisons of the means through a Brown-Forsythe and Welch ANOVA tests. The significance threshold was set at 0.05 for all tests.

## Supporting information

Supplementary references

## ACKNOWLEDGMENTS

We are deeply thankful to E. Bidnenko, V. Bidnenko, M. Boudvillain, S. Durand and C. Condon for fruitful discussions. We also thank Sandra Derozier and Cyprien Guérin for the visualisation of the transcriptomic data with Genoscapist.

