## Supplementary figures for "Quorum sensing dependent repression of the *Staphylococcus aureus* transcription termination factor Rho is necessary for pathogenicity"

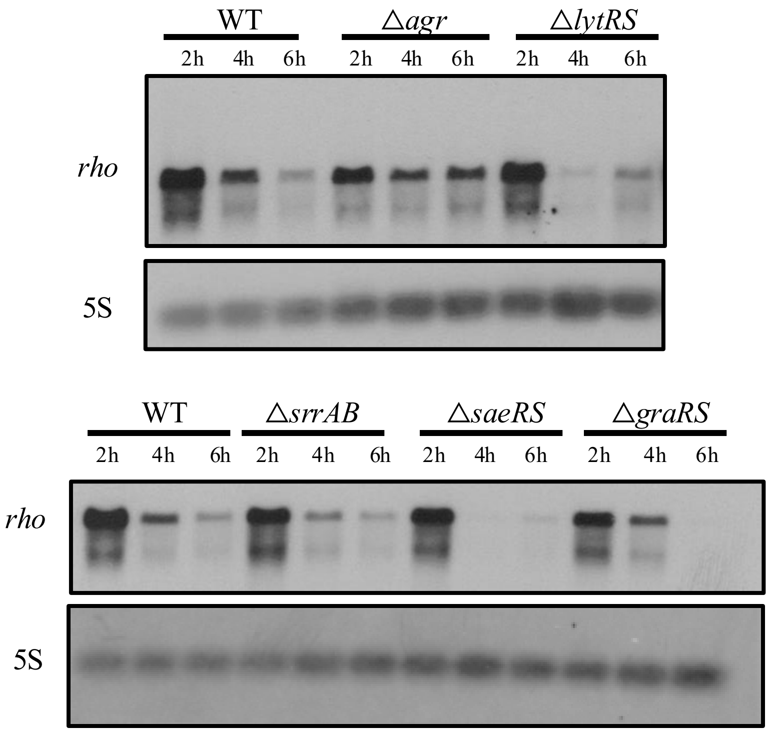


**Figure S1. Effect of the two-component systems on *rho* mRNA expression.**
Northern blot analysis of *rho* mRNA in HG001 WT and mutant strains carrying deletion of one two-component system. Total RNA was extracted from samples taken at 2, 4 and 6 h of growth in BHI medium. The loading control corresponded to 5S rRNA (5S) was probed on the same membrane.

**A.**

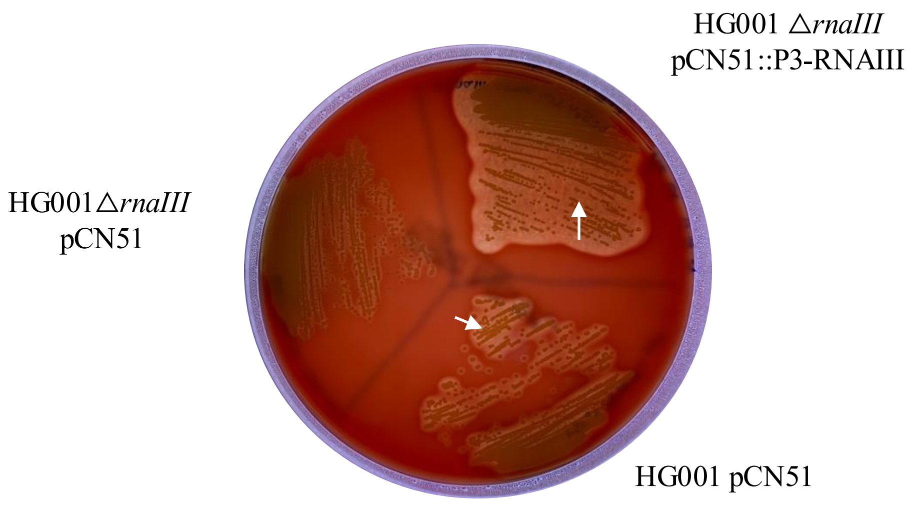


**B.**

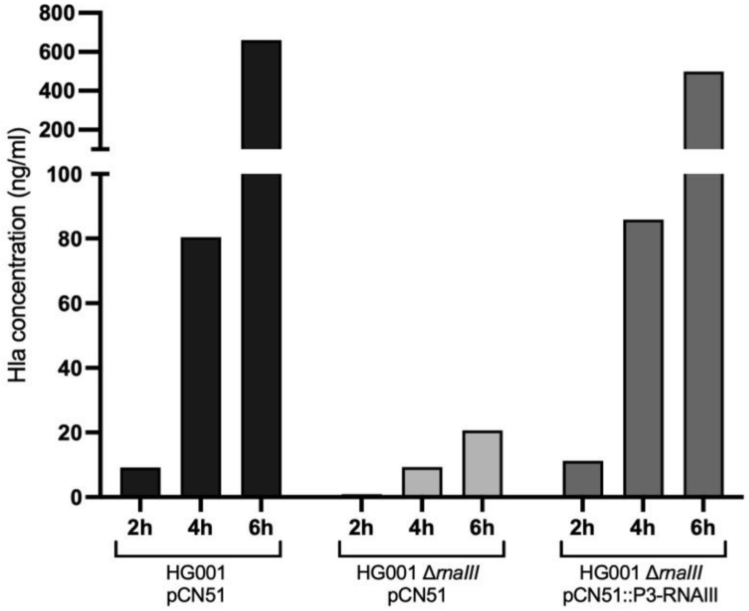


**Figure S2. Complementation of RNAIII restores hemolytic activity of Δ*rnaIII* mutant.**
**A.** HG001 pCN51, HG001 Δ*rnaIII* pCN51 and HG001 Δ*rnaIII* pCN51::P3-RNAIII strains grown on blood-agar medium. Hemolytic halos were observed for HG001 pCN51 and the complemented HG001Δ*rnaIII* pCN51::P3-RNAIII strains denoted by white arrows. **B**. ELISA analysis of *in vivo* expression of α-hemolysin in HG001 pCN51, HG001Δ*rnaIII* pCN51 and HG001Δ*rnaIII* pCN51::P3-RNAIII strains. Strains were grown in BHI medium with 10 µg/mL erythromycin and culture supernatants were taken at different time points (2 h, 4 h and 6 h). Hla quantification was performed by a sandwich ELISA using custom-made antibodies.

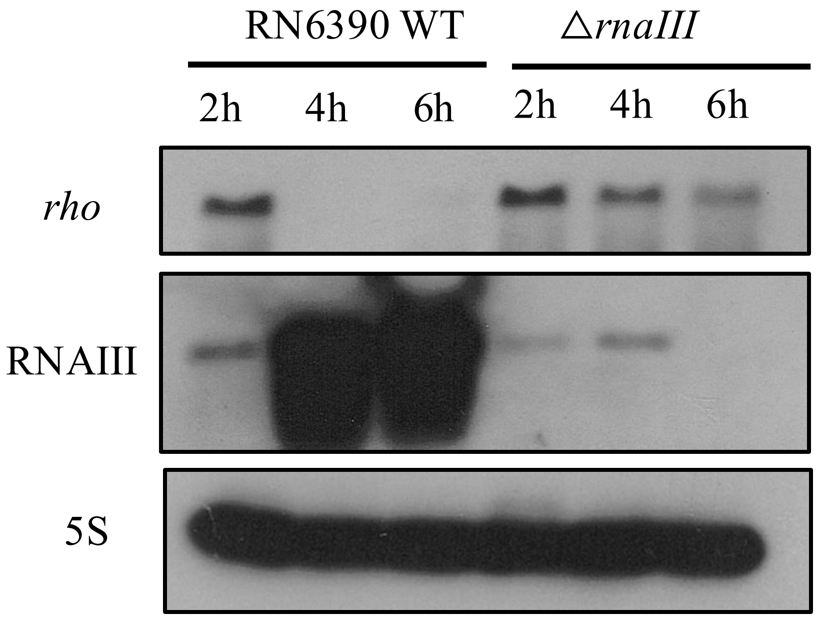


**Figure S3. Expression of *rho* mRNA is RNAIII-dependent in *S. aureus* RN6390.**
Northern blot analysis of *rho* mRNA in RN6390 WT and Δ*rnaIII* mutant strain (deletion of *rnaIII*). Total RNA was extracted at 2, 4 and 6 h of growth in BHI medium at 37°C and loaded on a 1% agarose gel. DIG-labeled RNA probes against *rho* mRNA, and the loading control 5S rRNA (5S) were used on the same membrane, while the same samples were run on a second gel in parallel to probe against RNAIII.

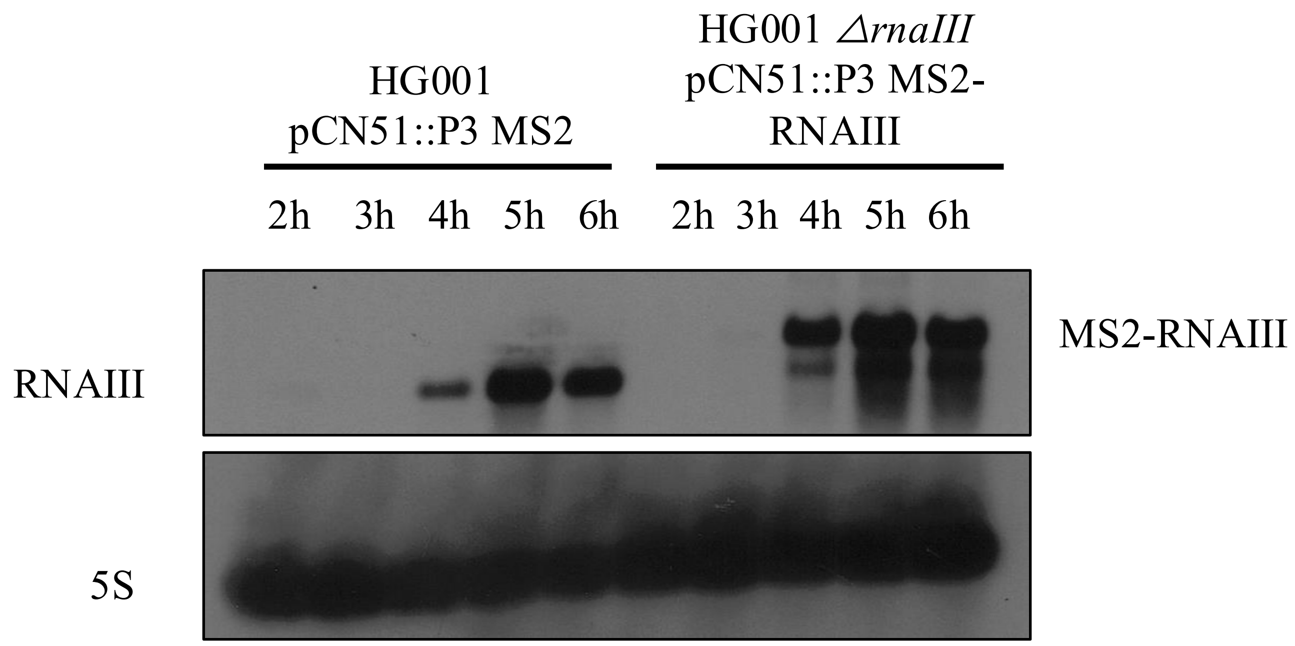


**Figure S4. Expression of RNAIII in strains used for MAPS.**

Northern blot analysis of RNAIII in HG001 pCN51::P3 MS2 expressing the MS2 tag alone and HG001 *ΔrnaIII* pCN51::P3 MS2-RNAIII expressing MS2-RNAIII. Cultures were grown in BHI medium at 37°C and samples were taken at different times. Total RNA was extracted and loaded on a 1% agarose gel to perform Northern blot and assess the expression of the MS2-tagged RNAIII. DIG-labeled RNA probes against RNAIII and the loading control 5S rRNA (5S) were used on the same membrane.

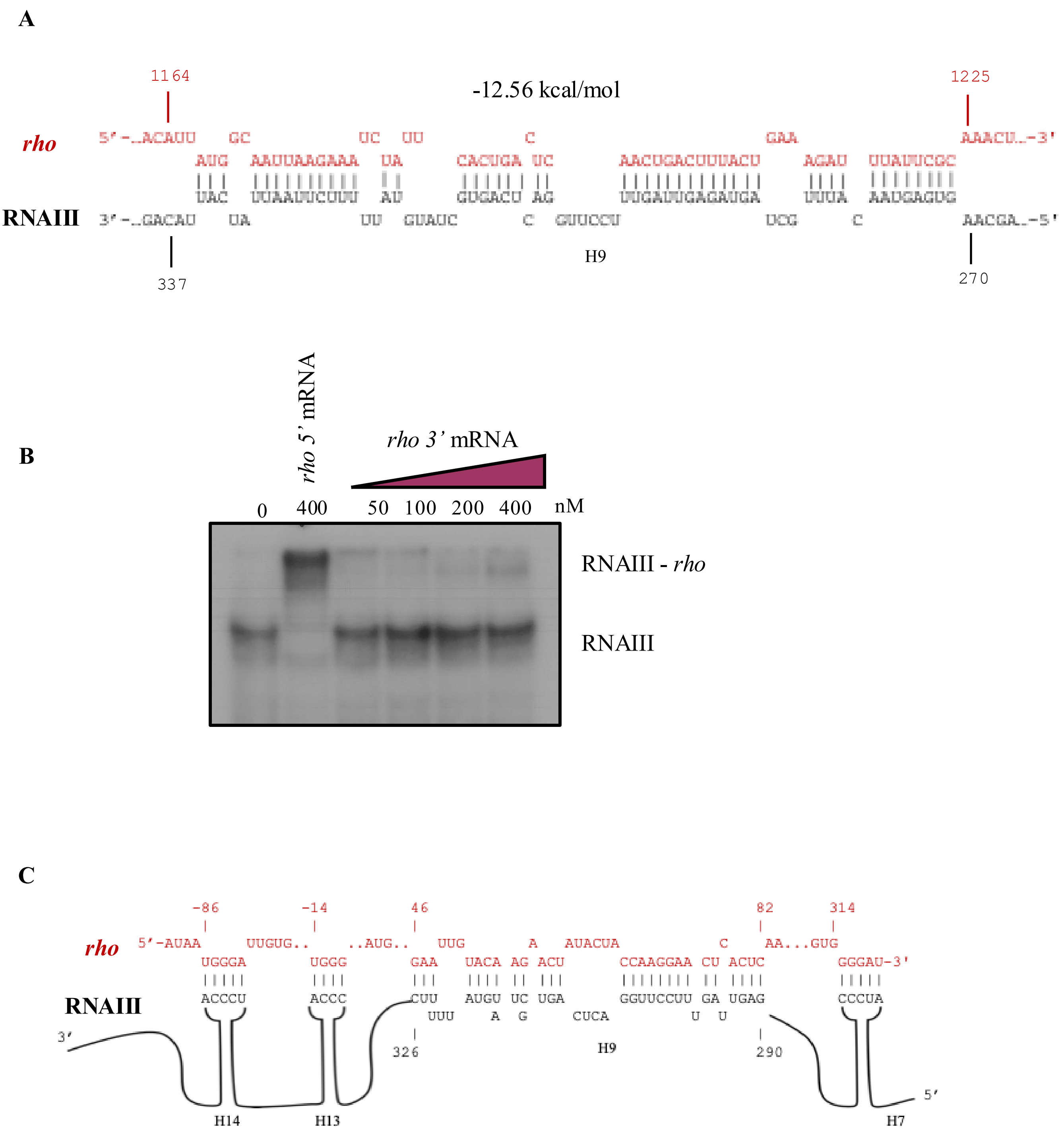


**Figure S5. The 3' region of *rho* mRNA does not significantly interact with RNAIII *in vitro*.**
**A.** Predicted interaction of the 3' region of the coding sequence of the *rho* mRNA with RNAIII using IntaRNA. **B.** Electrophoretic mobility shift assay (EMSA) of the 5'-end radiolabeled RNAIII incubated with either the 5' part (1-468 nts) of *rho* mRNA or with increasing concentrations of the 3' part (965-1417 nts) of *rho* mRNA. A control of 5' end labeled RNAIII alone is also shown. **C.** Proposed pairing scheme between RNAIII and *rho* mRNA. Several interaction sites include hairpins 7 (H7) and 9 (H9) and potentially hairpins 13 (H13) and 14 (H14) of RNAIII, and the 5'UTR and the coding sequence of *rho* mRNA.

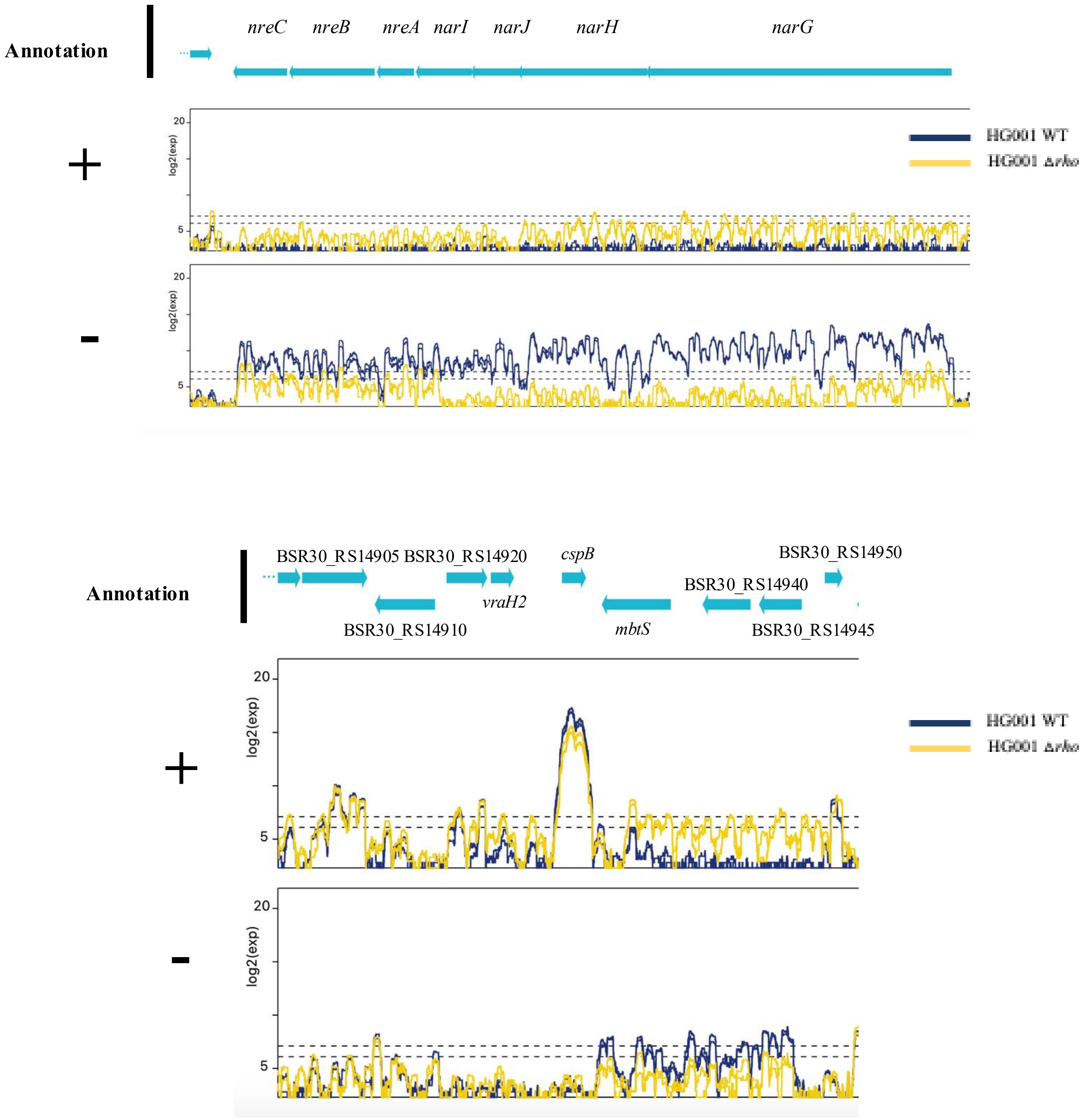


**Figure S6. Antisense RNAs generated upon *rho* deletion impact opposite strand mRNA expression.** Expression profiles of HG001 WT and HG001Δ*rho* strains measured by RNAseq during exponential phase of growth. Both strands (+ and -) are shown. The annotation of the genomic region is detailed at the top. Genes of unknown function (hypothetical genes and prophages) are shown with their reference number (BSR_RSXXXXX). Each strain's profile is represented with a different color, blue for WT and yellow for the Δ*rho* mutant. Decreased expression of the depicted regions are observed in the absence of *rho* with concomitant increase of an antisense RNA to the same region.

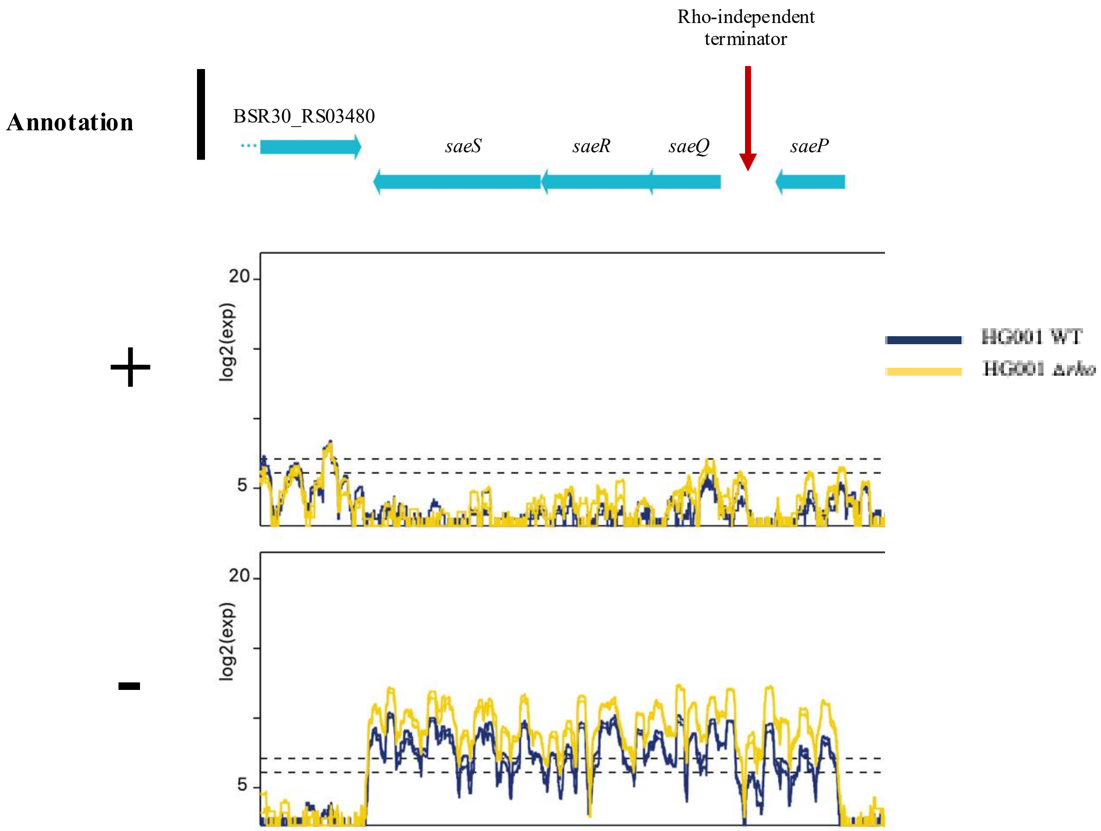


**Figure S7. The *saePQRS* operon is upregulated upon *rho* deletion.**
Expression profiles of HG001 WT and HG001Δ*rho* strains at the *saePQRS* locus measured by RNAseq during exponential phase of growth. Both strands (+ and -) are shown. The annotation of the genomic region is detailed at the top. Genes of unknown function (hypothetical genes and prophages) are shown with their reference number (BSR_RSXXXXX). Each strain's profile is represented with a different color, blue for WT and yellow for the Δ*rho* mutant. Rho-independent terminators are pointed with an arrow on the genomic locus.

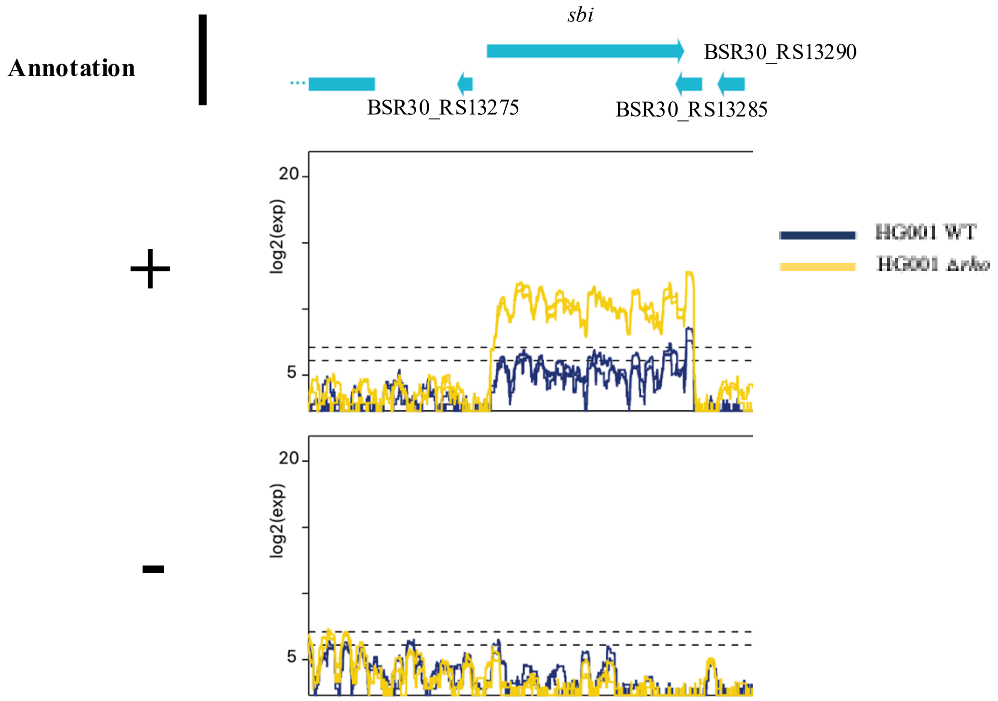


**Figure S8. Increased expression of the *sbi* mRNA in the Δ*rho* mutant.**
Expression profiles of HG001 WT and HG001Δ*rho* strains measured by RNAseq during exponential phase of growth. Both strands (+ and -) are shown. The annotation of the genomic region is detailed at the top. Genes of unknown function (hypothetical genes and prophages) are shown with their reference number (BSR_RSXXXXX). Each strain's profile is represented with a different color, blue for WT and yellow for the *rho* mutant.

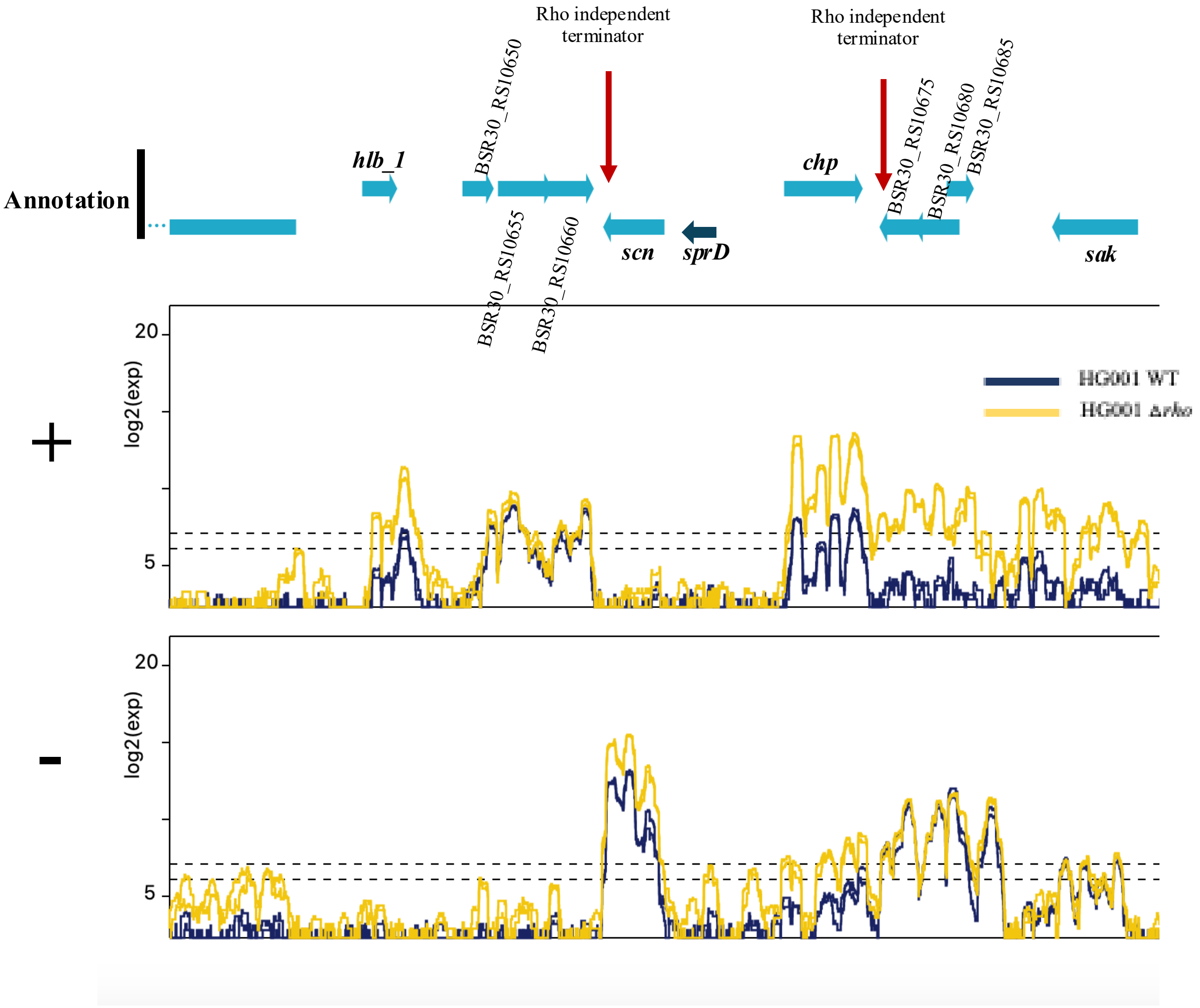


**Figure S9. Direct Rho-mediated control virulence genes.**

Expression profiles of HG001 WT and HG001Δ*rho* strains measured by RNAseq during exponential phase of growth. Both strands (+ and -) are shown. The annotation of the genomic region is detailed at the top. Genes of unknown function (hypothetical genes and prophages) are shown with their reference number (BSR_RSXXXXX). Each strain's profile is represented with a different color, blue for WT and yellow for the Δ*rho* mutant. Rho-independent terminators are pointed with an arrow on the genomic locus.

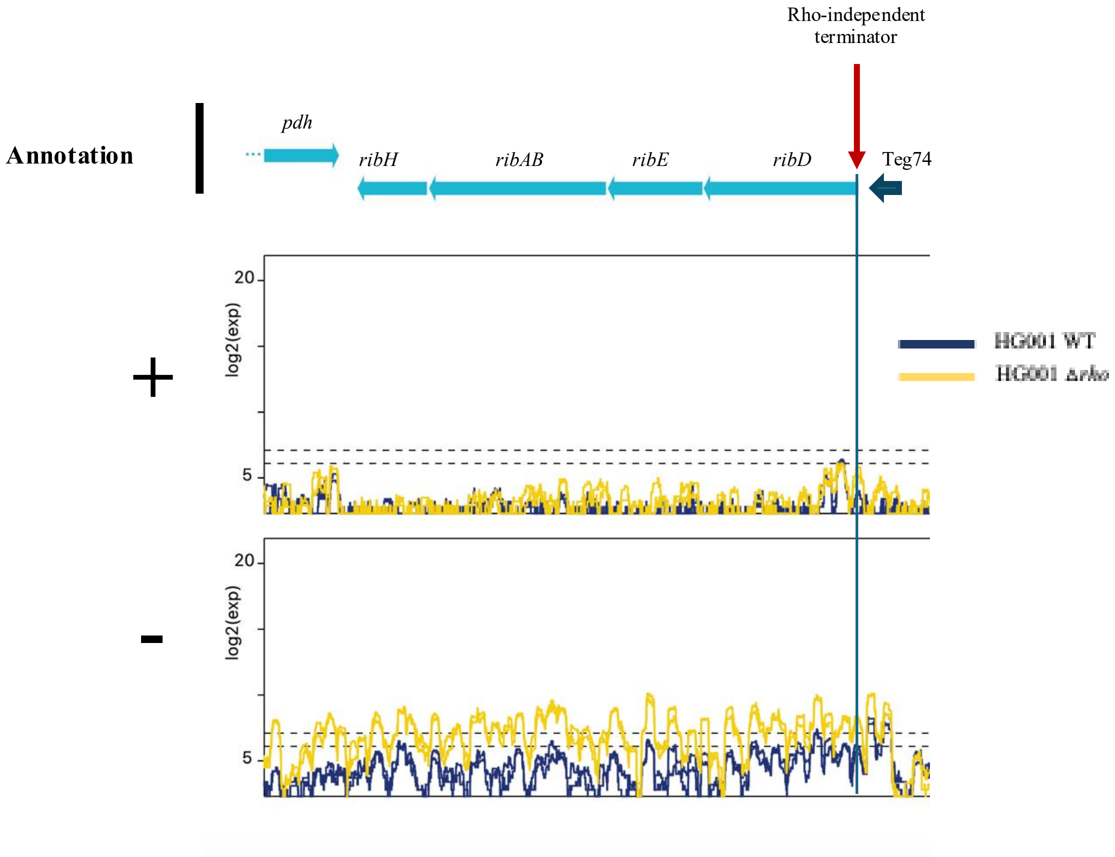


**Figure S10. Rho-mediated control of the FMN riboswitch.**
Expression profiles of HG001 WT and HG001Δ*rho* strains measured by RNAseq during exponential phase of growth. Both strands (+ and -) are shown. The annotation of the genomic region is detailed at the top. Genes of unknown function (hypothetical genes and prophages) are shown with their reference number (BSR_RSXXXXX). Each strain's profile is represented with a different color, blue for WT and yellow for the Δ*rho* mutant. Rho-independent terminators are pointed with an arrow on the genomic locus.

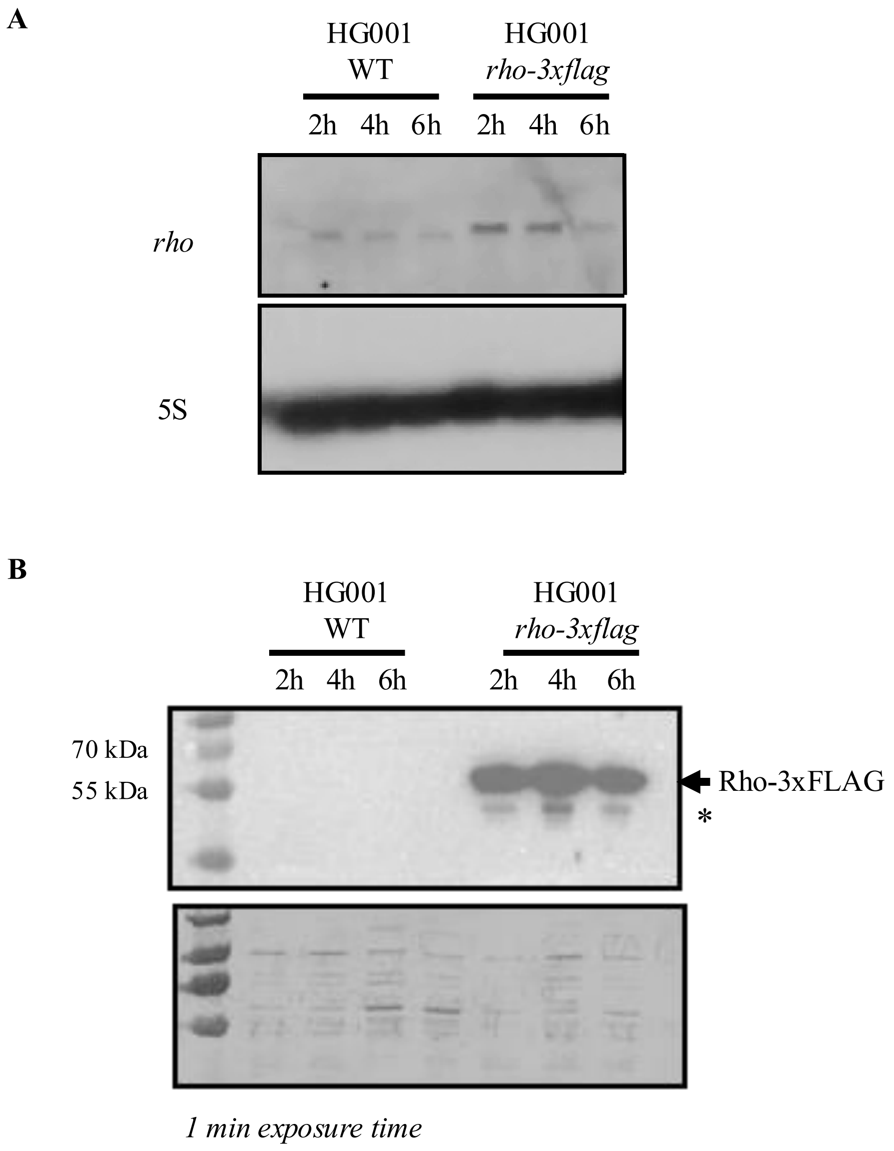


**Figure S11. Expression of Rho3x-FLAG.**
**A**. Northern blot analysis of *rho* in HG001 WT, HG001 *rho-3xflag.* Total RNA was extracted at 2, 4 and 6 h of growth in BHI medium at 37ºC and loaded on a 1% agarose gel. DIG-labeled RNA probes against *rho* and the loading control 5S rRNA (5S) were used on the same membrane. **B**. HG001 WT and HG001 *rho-3xflag* strains were grown in BHI medium at 37º C and samples were taken at different times. Cells were harvested, lysed and the supernatant was used for measuring total protein concentration. Samples of 10 µg of total proteins were loaded on a 12% polyacrylamide-SDS gel to perform Western blot. Membranes were incubated with anti-FLAG mouse antibodies followed by the incubation with a goat anti-mouse peroxidase (HRP) antibody. A protein extract from HG001 Δ*rho* was also loaded as a control. Two exposition times are shown. *=non-specific signal.

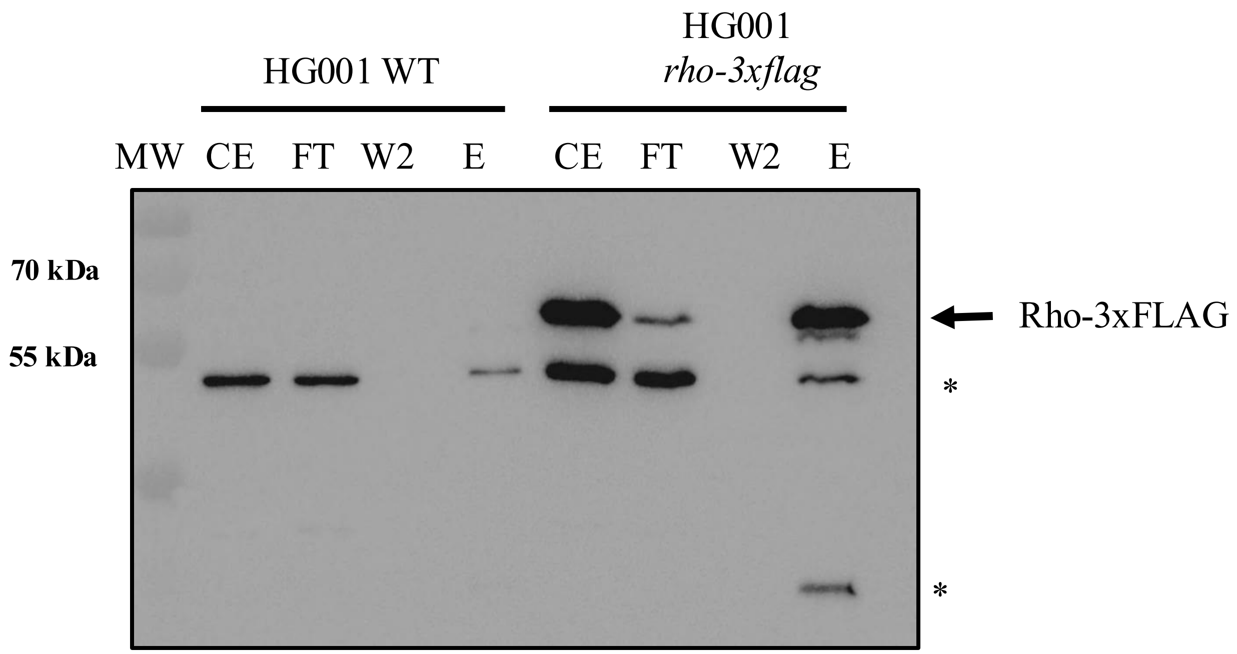


**Figure S12. Immunoprecipitation of Rho 3x-FLAG.**
HG001 WT and HG001 *rho-3xflag* strains were grown in duplicated in BHI medium at 37ºC until exponential phase. Cells were harvested, lysed and supernatants were used for immunoprecipitation using anti-FLAG M2 affinity gel. Samples from each fraction (CE: crude extract, F: flowthrough, W2: wash 2 and E: elution) were collected and loaded on a 12% polyacrylamide-SDS gel to perform Western blot. Membranes were incubated with anti-FLAG mouse antibodies followed by the incubation with a goat anti-mouse peroxidase (HRP) antibody. The recombinant Rho-3xFLAG protein was successfully recovered after elution. RNAs from this fraction were isolated and sequenced. *=non-specific signal. The experiment was reproduced three times.

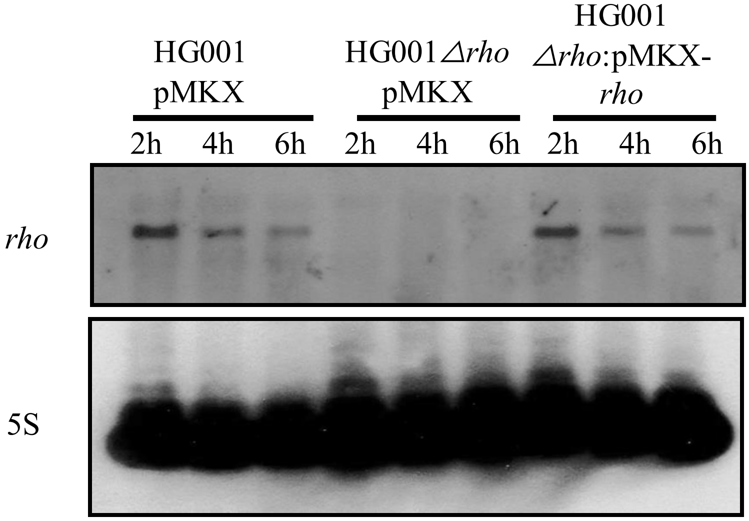


**Figure S13. Leaky expression of *rho* mRNA from a plasmid results in WT *rho* mRNA levels.** Northern blot analysis of *rho* mRNA in HG001 pMKX, HG001*Δrho* pMKX and HG001*Δrho* pMKX-*rho* strains. Total RNA was extracted at 2, 4 and 6 h of growth in BHI medium at 37°C without xylose induction and loaded on a 1% agarose gel. DIG-labeled RNA probes against *rho* and the loading control 5S rRNA (5S) were used on the same membrane. Expression of *rho* follows the same pattern as WT *rho* regulation in the complemented strain without xylose induction.

| **Strains** | **Culture 1** | **Culture 2** | **Culture 3** | **Culture 4** |
|---|---|---|---|---|
| HG001 pMKX | 32 | 16 | 16 | 16 |
| HG001Δ*rho* pMKKX | 128 | 64 | 128 | 64 |
| HG001Δ*rho* pMKKX-*rho* | 32 | 32 | 32 | 32 |

**Figure S14: Quantification of coagulase activity in various strains.**
The coagulase tube test was performed with four supernatants from HG001 WT strains, HG001Δ*rho* mutant strain and HG001Δ*rho* pMKK-*rho* complemented with a plasmid expressing *rho*. Cultures were grown during 5 h in BHI (n=4). These supernatants were diluted in serial two-fold dilution and mixed with rabbit plasma. The titer was the reciprocal of the highest dilution of the supernatant that showed evidence of clotting after incubation for 18 h at 37°C.
